# Insights into the early-life chemical exposome of Nigerian infants and potential correlations with the developing gut microbiome

**DOI:** 10.1101/2023.11.08.566030

**Authors:** Ian Oesterle, Kolawole I. Ayeni, Chibundu N. Ezekiel, David Berry, Annette Rompel, Benedikt Warth

## Abstract

Early-life exposure to natural and synthetic chemicals can impact acute and chronic health conditions. Here, a suspect screening workflow anchored on high-resolution mass spectrometry was applied to elucidate xenobiotics in breast milk and matching stool samples collected from Nigerian mother-infant pairs (n = 11) at three time points. Potential correlations between xenobiotic exposure and the developing gut microbiome, as determined by 16S rRNA gene amplicon sequencing, were subsequently explored. Overall, 12,192 and 16,461 features were acquired in the breast milk and stool samples, respectively. Following quality control and suspect screening, 562 and 864 features remained, respectively, with 149 of these features present in both matrices. Taking advantage of 242 authentic reference standards measured for confirmatory purposes of food bio-actives and toxicants, 34 features in breast milk and 68 features in stool were identified and semi-quantified. Moreover, 51 and 78 features were annotated with spectral library matching, as well as 416 and 652 by *in silico* fragmentation tools in breast milk and stool, respectively. The analytical workflow proved its versatility to simultaneously determine a diverse panel of chemical classes including mycotoxins, endocrine-disrupting chemicals (EDCs), antibiotics, plasticizers, perfluorinated alkylated substances (PFAS), and pesticides although it was originally optimized for polyphenols. Spearman rank correlation of the identified features revealed significant correlations between chemicals of the same classification such as polyphenols. One-way ANOVA and differential abundance analysis of the data obtained from stool samples revealed that molecules of plant-based origin were elevated when complementary foods were introduced to the infants’ diets. Annotated compounds in the stool, such as tricetin, positively correlated with the genus *Blautia*. Moreover, vulgaxanthin negatively correlated with *Escherichia*-*Shigella*. Despite the limited sample size, this exploratory study provides high-quality exposure data of matched biospecimens obtained from mother-infant pairs in sub-Saharan Africa and shows potential correlations between the chemical exposome and the gut microbiome.

**Highlights:**

- Suspect screening of exposure biomarkers in human breast milk and infant stool.
- 542 features in breast milk and 864 in stool were identified or annotated.
- Consumption of complementary foods influenced the chemical exposure of infants.
- Correlations between xenobiotics in both biological matrices evaluated.
- Dietary exposure correlated to the stool microbiome composition.

**Graphical abstract:** 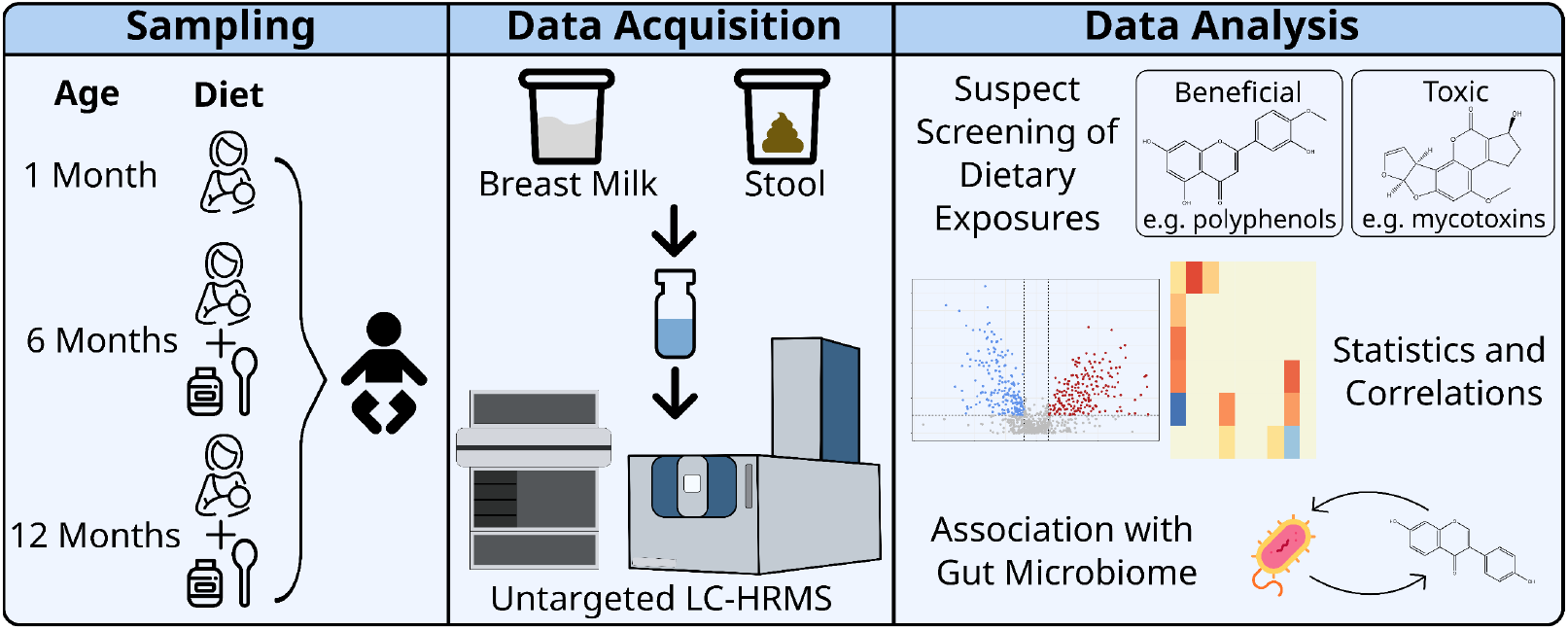

## 1. Introduction

In 2005, Christopher Wild coined the term “exposome”, a currently expanding research field that investigates the totality of chemicals that an individual is exposed to during their lifetime, and related health consequences (Miller and Jones 2014). It focuses often on co-exposure and toxicological mixture effects of xenobiotics and their biotransformation products and can be further complemented with other -omic techniques, e.g. metagenomics, to characterize the biological impact of these exogenous chemicals (Vitale et al. 2021, Kalia et al. 2022). Moreover, emphasis is given to the first thousand days of life, as exposure during this period is suspected to especially impact the development of chronic diseases later in life (Heindel et al. 2015). For example, exposure to endocrine disrupting chemicals (EDCs) has been associated with an increased risk of diseases like neurodevelopmental disorders (Braun 2017). A major source of exposure during early-life is the infant’s diet. For instance, consumption of complementary foods rather than breast milk can expose infants to higher levels of mycotoxins (Krausová et al. 2023).

Exposome research has mainly focused on toxic exposures and often does not include molecules that might be able to mitigate adverse actions, e.g. polyphenols which are prevalent in plant-based foods. Polyphenols exhibit various properties in humans that are typically described as positive, like antioxidant and antimicrobial effects (Pandey and Rizvi 2009), or potentially negative, such as estrogenic effects (Křížová 2019). Moreover, polyphenols can reduce the toxicity of other xenobiotics, as described for mycotoxins (Rasouli et al. 2022).

Environmental exposure (Claus et al. 2016) and diet (David et al. 2014, Wilson et al. 2020) can also modulate the gut microbiome, which can further impact human health (Hou et al. 2022). Conversely, the microbiome can influence xenobiotics (Collins and Patterson 2020), leading to a bi-directional relationship (Clarke et al. 2019). Thus, xenobiotics can contribute to dysbiosis during early-life (Ayeni et al. 2022), especially as the microbiome of infants is not yet fully established (Wopereis et al. 2014). Therefore, understanding xenobiotic and early-life microbiome interactions is vital.

As xenobiotics are chemically diverse, assessing human exposure requires cutting-edge exposomic workflows anchored on mass spectrometry (MS) (Bocato et al. 2019, Flasch et al. 2022b, Gu et al. 2023). Untargeted approaches, compared to targeted approaches, are advantageous as they allow the exploration of a broader spectrum of analytes (Oesterle et al. 2021). However, only features, defined as a detected signal intensity and an associated *m/z* and retention time value, are acquired in untargeted approaches. The annotation of analytes is challenging and supported by fragmentation spectra. One method to acquire features and their fragmentation spectra simultaneously is sequential window acquisition theoretical hold (SWATH) data-independent acquisition (DIA), which involves fragmenting all ions in sequential windows of monoisotopic masses (Bonner and Hopfgartner 2019). Though data analysis is more complex than data-dependent acquisition (DDA), SWATH DIA yields fragmentation spectra for all features present (Guo and Huan 2020), allowing better retrospective annotation. As untargeted approaches generate a high number of features, suspect screening can be employed to filter for analytes of interest (Pourchet et al. 2020).

Several studies have reported on metabolomic profiles of healthy children (Chen et al. 2019, Holzhausen et al. 2023), children with severe acute malnutrition (McMillan et al. 2016), or comparing breastfed infants with infants fed formula (Brink et al. 2020, Silner et al. 2021). However, to our knowledge, there is limited data on the longitudinal exposomic/metabolomic profiles of neonates and infants via breast milk and matched stool samples. Thus, the objectives of this study were to apply a recently developed untargeted exposomic workflow (Oesterle et al. 2023) utilizing SWATH DIA on breast milk and stool collected longitudinally from Nigerian mother-infant pairs. This workflow allowed to 1) elucidate exposure profiles of potentially beneficial and toxic xenobiotics present, 2) investigate changes in xenobiotic profiles as complementary foods were introduced in the diet, and 3) correlate xenobiotics detected in the infants’ stool to their gut microbiome. This research therefore aims to give preliminary insights into the chemical exposome during a key age window of a high exposure population.

## 2. Materials and methods

### 2.1. Study design and ethical approval

The longitudinal pilot study involved human breast milk and matched infant stool samples from Ilishan-Remo, Ogun state, Nigeria. Details of study location and sample collection were previously described by Ayeni et al. (2024). The same set of samples was used before to investigate specific exposure classes: polyphenols in the breast milk samples (Berger et al. 2024), and mycotoxins in the stool (Krausová et al. 2022) and breast milk (Ayeni et al. 2024). Ethical approval was obtained from the Ethical Committee of Babcock University (BUHREC 421/21R, BUHREC 466/23). All mothers were properly informed before providing their written consent to be included in the study.

### 2.2. Reagents and chemicals

In total, 242 authentic reference standards of different xenobiotic classes were used to allow for Level 1 identifications, benchmarking the approach, and enabling semi-quantification. Stock solutions were prepared by diluting the standards in either methanol (polyphenols) or acetonitrile (other standards) and used to prepare working mixes for spiking experiments. Information on all standards and additional reagents are listed in Table A.1.

### 2.3. Sample preparation

The breast milk samples were prepared following the protocol optimized by Berger et al. (2024). The infant stool samples were prepared following the method of Krausová et al. (2022) with minor modifications as explained in Supplementary Information (SI) B. The quantities of wet stool and dry stool for each sample are listed in Table A.2.

### 2.4. LC-MS instrumentation

An ultra-high-performance LC (UHPLC)-electrospray ionization (ESI)-quadrupole time-of-flight (QTOF)-high-resolution MS (HRMS) UHPLC-ESI-QTOF-HRMS system consisting of an Agilent 1290 Infinity II UHPLC and a *Sciex* ZenoTOF 7600 MS was used to analyze the samples. The LC parameters were previously optimized (Oesterle et al. 2022) and are listed in SI Table A.3. The MS was operated in SWATH mode with the windows optimized for each matrix individually using the total ion chromatogram from a pooled quality control sample. MS parameters are reported in Table A.4.

### 2.5. LC-MS data processing

The acquired data was processed with MS-Dial (v4.9.221218) (Tsugawa et al. 2015). MS-Finder (v3.52) (Tsugawa et al. 2016) was used for *in silico* fragmentation. The identification levels of the feature annotation were given based on the levels previously defined by Schymanski et al. (2014), with the modification of Level 3 as described by Oesterle et al. (2023). The detailed data processing procedure is explained in SI B and the parameters are reported in Table A.4.

### 2.6. LC-MS Quality control

Pooled quality control (QC) samples were prepared for each biological matrix. The QC samples were then analyzed throughout the corresponding sequence. In addition, a process blank was prepared for each sample preparation procedure. The QC and process blank samples were used for monitoring the sequence and correcting the data processing (removing features arising from noise and creating matrix-matched calibration curves). The detailed quality control procedures are reported in SI B.

### 2.7. 16S ribosomal RNA (rRNA) gene amplicon sequencing of the infant stool

Gut microbiome data of the infants’ stool samples obtained at months 1, 6 and 12 post-delivery were retrieved from Ayeni et al. (2024). The detailed procedure of DNA extraction, polymerase chain reaction, and 16S rRNA gene amplicon sequencing applied to the infants’ stool was previously published (Pjevac et al. 2021).

### 2.8. Statistical analysis

Following data processing, the chromatographic peak areas of the stool features were normalized by their sample dry weight (Table A.2). MetaboAnalyst (v5.0) (Pang et al. 2022) was then used to normalize the distribution of the features and for their statistical analysis. ChemRICH (v4.0) (Barupal and Fiehn 2017) was used to generate chemical enrichment plots with the results from MetaboAnalyst. In addition, R (v4.3.1) (R Core Team 2023) was used for further statistical analysis of the features and microbiome data. Further details of the statistical analysis are explained in SI B.

## 3. Results and discussion

### 3.1. Suspect screening of breast milk and stool samples

Overall, a total of 12,192 features were acquired in the breast milk and 16,461 features in the stool samples. After quality control, 4,347 and 6,905 features remained, respectively. A suspect screening workflow optimized for polyphenols but also capable of covering many other xenobiotics and endogenous metabolites (Oesterle et al. 2023) was used to extract features of interest. The application of this workflow resulted in a total of 542 matched features in the breast milk and 864 in the stool samples. From the 542 breast milk features (Table A.6), 34 were identified as Level 1, 51 were annotated as Level 2a, 416 as Level 3a, and 41 as Level 3b. From the 864 features in the stool (Table A.7), 68 were identified as Level 1, 78 were annotated as Level 2a, 652 as Level 3a, and 66 were putatively annotated as Level 3b.

For Level 1 identification, 242 authentic reference standards representing diverse xenobiotics were utilized. These standards included air pollutants, disinfection by-products, endogenous estrogens, food processing by-products, industrial side-products, pesticides, mycotoxins, perfluorinated alkylated substances, personal care product/pharmaceuticals, phytotoxins, plasticizer / plastic components, antibiotics, and polyphenols (Table A.1). The Level 1 identified features were also semi-quantified (Table A.8 and SI B), and their concentrations are represented as boxplots (Figures 1 and B.3 for breast milk, and Figures 2 and B.4 for stool).

In breast milk, many of the features detected were fatty acids, peptides, or amino acids (Figure 1a). This was expected as breast milk is rich in nutrients, especially lipids, proteins, and carbohydrates, required for infant growth and development (Boudry et al. 2021). A heatmap of the annotated features (Figure 1b) depicts that there are two main clusters, though there is no clear separation of the types of features in each of the two groups. Moreover, no clear pattern in the concentration of the identified features at the three time points can be observed (Figure 1c). In addition to nutrients, a variety of polyphenols were detected in the breast milk. For instance, daidzein was detected at concentration ranges between 0.0035 to 12 ng/mL (Table A.8, Figure 1c), which was similar to the reported concentrations from a longitudinal study of an Austrian mother (Jamnik et al. 2022). The breast milk samples from this cohort were previously analyzed with a targeted LC-MS/MS workflow focusing on polyphenols (Berger et al. 2024), and similar analytes and patterns were observed between both assays. Several studies have shown that polyphenols are abundant in human breast milk and they could be beneficial to infants (Lu et al. 2021, Song et al. 2013, Carregosa et al. 2023).

**Figure 1.**
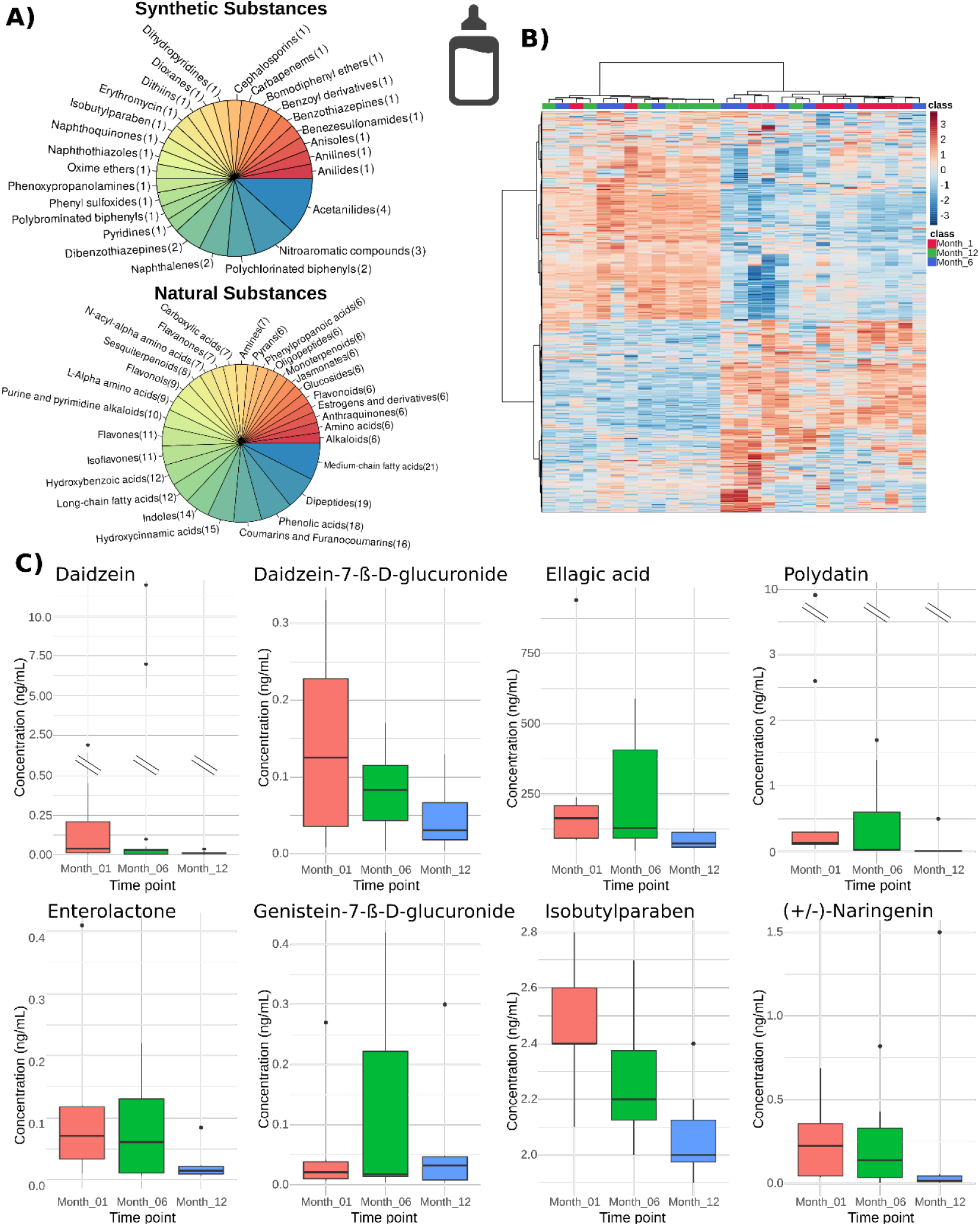
**A)** The major chemical classes of the annotated features in the breast milk samples at three different time points represented in two pie charts: one with synthetic substances and one with endogenous and exogenous natural substances. **B)** Heatmap with Ward clustering of all the features detected in the breast milk samples. **C)** Box plots of selected semi-quantified features (Table A.8) over time that represent both potentially beneficial and toxic xenobiotics in the breast milk samples. The boxplots of all the identified features are shown in Figure B.2.

Many toxicants such as mycotoxins are found at lower prevalence and concentrations in breast milk (Adejumo et al. 2013, Braun et al. 2020). For example, only two potentially toxic xenobiotics, alternariol monomethyl ether and isobutylparaben, were identified at Level 1. Alternariol monomethyl ether, which ionizes highly efficient in ESI and thus typically has a very low limit of detection, was previously reported as common in breast milk from a Nigerian cohort (Braun et al. 2022, Ezekiel et al. 2022). While there is scarce reported data on parabens in breast milk from Nigeria, these xenobiotics have been reported in breast milk from other regions (Fisher et al. 2017, Kim et al. 2023). The antibiotic, azithromycin, was detected in only one sample at a concentration of 1100 ng/mL (Table A.8). Interestingly, the participant who donated this sample used azithromycin for medical treatment purposes (unpublished data), and it was detected in the corresponding infant stool sample (Table A.8). This observation suggests that azithromycin can be transferred from mother’s breast milk to infant (Roca et al. 2015).

More diverse chemical classes, predominantly of plant-based origin, were present in the stool (Figure 2a). Moreover, many of the features were conjugated with sugar moieties. This could be attributed to phase II metabolism, as it was previously reported that metabolites of genistein included hexose and pentose conjugates in cells (Flasch et al. 2022a); or to their low intestinal absorption, as, for example, seen with the bioavailability of large polyphenols like proanthocyanidins (Scalbert et al. 2002). In the heatmap of the features detected in the stool (Figure 2b), two clusters are formed. One cluster was from the samples during the time of breastfeeding (month 1) and the other for those acquired after the introduction of complementary foods (months 6 and 12). In the quantified features, a similar pattern could be observed over time. Features belonging to various classes, such as catechins, flavanones, and mycotoxins, increased in concentration from month 1 to 12. Moreover, the concentrations were substantially higher at month 6 compared to month 1, when complementary foods were introduced to the infants by month 6 (Figure B.1). This observation suggests that complementary foods have a considerable influence on the features in stool. In addition, the presence of various features when the infants were one month old showed that bioactive compounds are transferred from the breast milk to the infants. For instance, one analyte that is most likely transferred via breast milk due to its stable concentrations over time (Figure B.4) is urolithin A, a microbial metabolite that showed properties beneficial to health (D’Amico et al. 2021).

**Figure 2.**
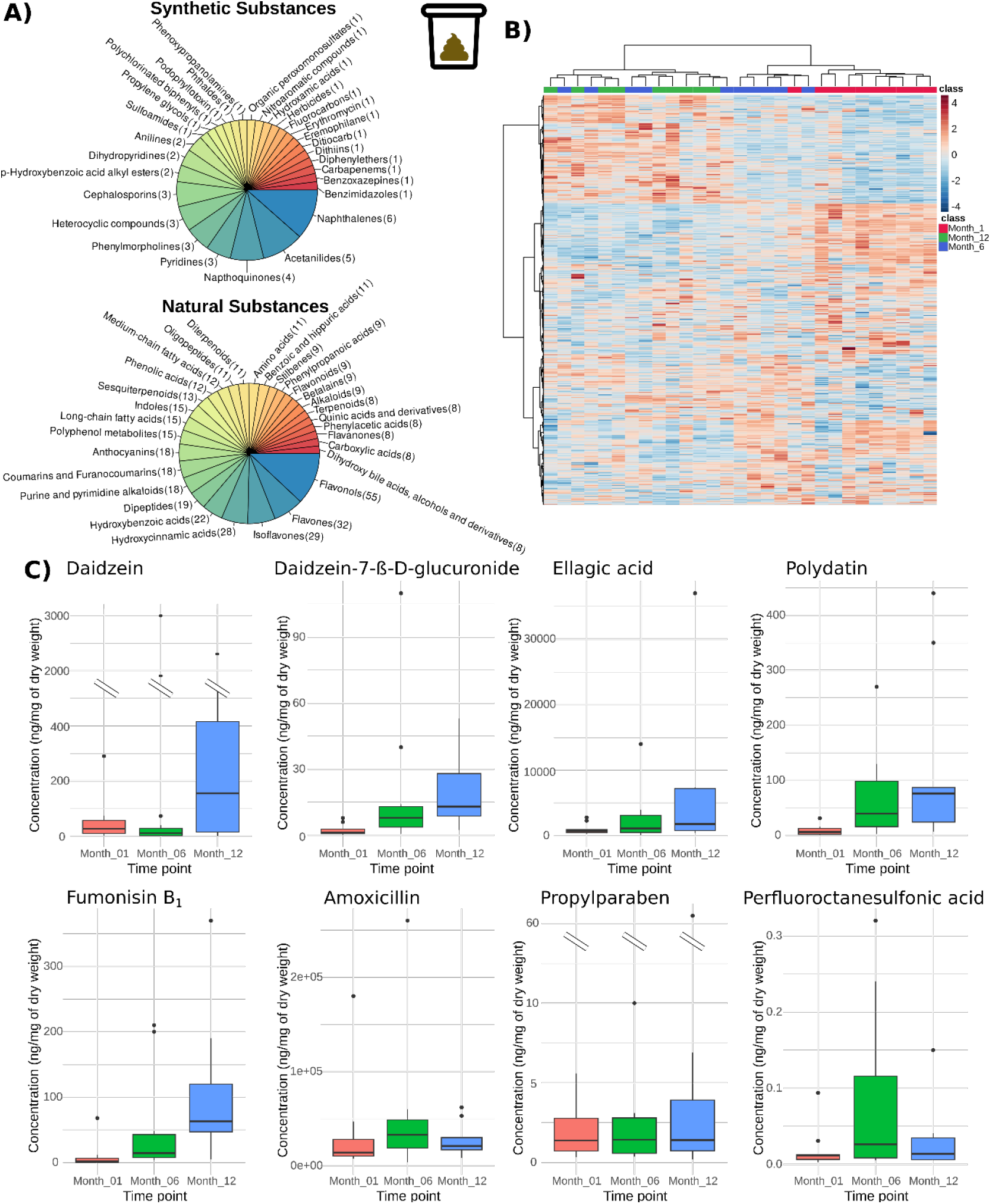
**A)** The major chemical classes of the annotated features in the infant stool at three different time points represented in two pie charts: one with synthetic substances and one with endogenous and exogenous natural substances. **B)** Heatmap with Ward clustering of all the features detected in the stool samples. **C)** Box plots of several semi-quantified features (Table A.8) at the three distinct infant ages. These represent both potentially beneficial and toxic xenobiotics and show that complementary foods increase exposure to various xenobiotics, while breast feeding keeps exposure levels low, despite potential lactational transfer. The box plots of all the identified features are shown in Figure B.3.

As the LC-MS method was originally optimized for polyphenols (Oesterle et al. 2022, Oesterle et al. 2023), it was expected that they would be one of the main chemical classes detected. However, a variety of other classes were determined, including several potentially toxic xenobiotics. For example, two mycotoxins, fumonisin B_1_ (Figure 3a) and nivalenol, were detected and quantified. Fumonisin B_1_ has been previously associated with neural tube defects and the etiology of esophageal cancer in humans (Marasas et al. 2004, Missmer et al. 2006). Moreover, this mycotoxin was previously found in stool of the same cohort using a targeted LC-MS/MS-based assay (Krausová et al. 2022, Ayeni et al. 2024), and the longitudinal pattern of occurrence was comparable (Ayeni et al. 2024). This highlights that targeted and non-targeted LC-MS/MS approaches can be complementary (Flasch et al. 2023). Perfluorooctanesulfonic acid, a toxic perfluorinated alkylated substance, was also detected (Figure 2c and 3b). For all infants, except one, the concentrations were lower during exclusive breastfeeding (month 1 to month 6) compared to when complementary foods were introduced (months 6 to 12). Moreover, perfluorooctanesulfonic acid was detected in Ogun river (>10 ng/L) (Ololade et al. 2018), a river in the region where the samples were collected. While many of the identified toxicants came from dietary sources, others may be attributed to other routes of exposure such as air, water, or cosmetics. Several antibiotics were also detected (Figure B.4), such as amoxicillin and oxytetracycline, and their presence can be attributed to the infants taking these antibiotics for medical treatment, or transferred via breastmilk, as with azithromycin. A variety of fatty acids were also detected in the stool, including arachidonic acid (Level 2a) and eicosapentaenoic acid (Level 3a). The levels of these two fatty acids were previously reported to be higher in stool of infants that were breastfed compared to infants fed that were formula-fed (Sillner et al. 2021).

**Figure 3.**
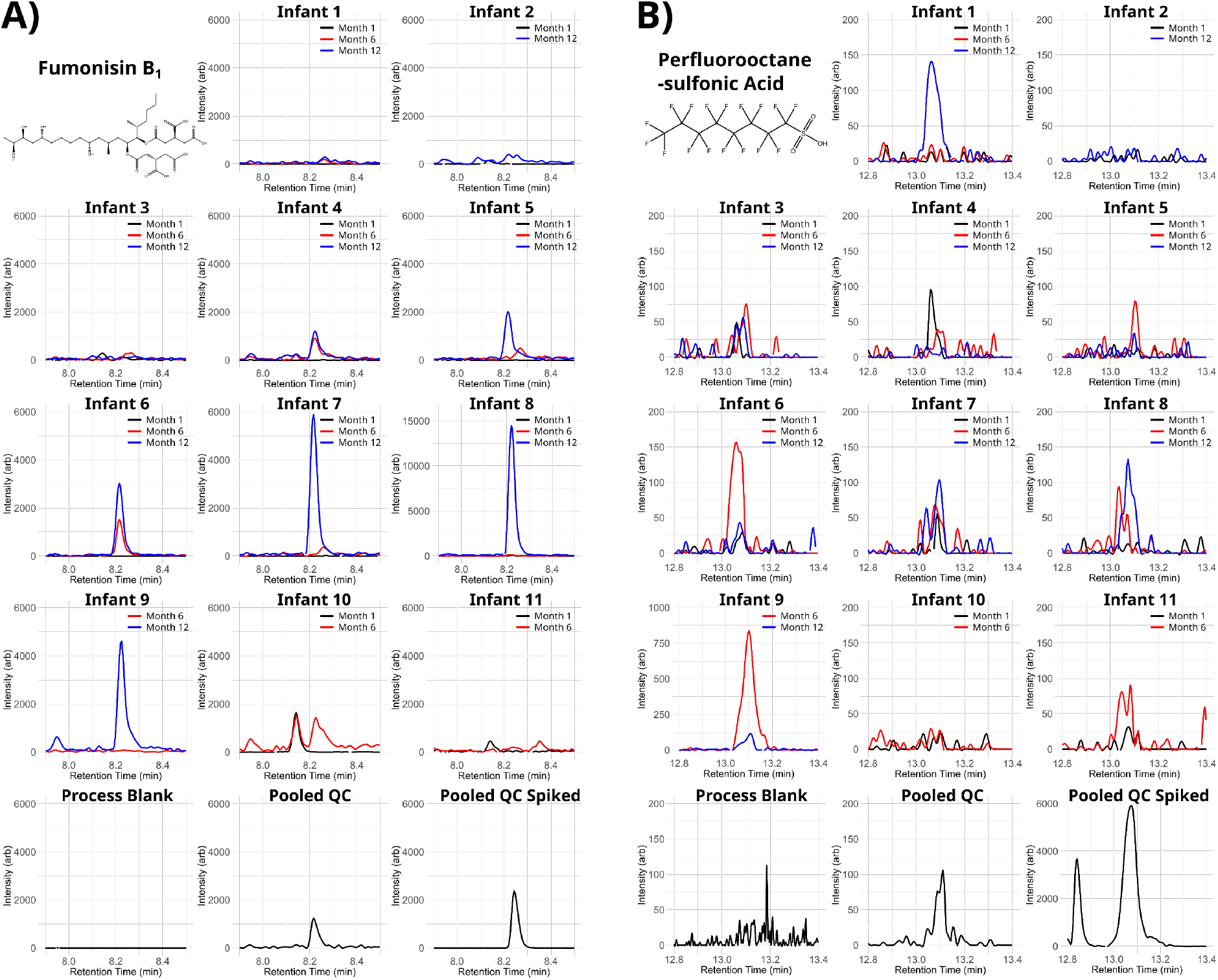
Extracted ion chromatograms of fumonisin B_1_ (**A**) and perfluorooctanesulfonic acid (**B**) in all infants at the three distinct time points. In addition, the process blank, pooled QC, and pooled QC spiked with the authentic reference standards (6.3 ng/mL for fumonisin B1 and 0.38 ng/mL for perfluorooctanesulfonic acid) are depicted. These chromatograms illustrate that exposure levels of adverse xenobiotics during exclusive breastfeeding were typically low and an increase was observed as complementary foods were introduced.

The features from each biological matrix were then compared with each other with a retention time deviation of 0.1 min and a mass error of 20 ppm. This resulted in only 149 of the 1,259 total features that were detected in both matrices, out of which 32, 24, 91, and four were annotated as Levels 1, 2a, 3a, and 3b, respectively. The high contrast between the two matrices is seen in the heatmap (Figure B.2) of all the features detected in both matrices. One of the most prevalent chemical classes in both biological matrices was phenolic acids, which is a class that comprises of many human and microbial metabolites (de Ferrars et al. 2014). Several other biotransformation products were detected in both matrices including caffeic acid-3-ß-D-glucuronide, daidzein-7-ß-D-glucuronide, and genistein-7-ß-D-glucuronide. This observation suggests that the chemicals were either transferred directly from the breast milk to the infant and the corresponding stool samples or that the infant’s metabolism conjugated the parent compounds in the colon or liver by UDP-glucuronosyltransferases (Rowland et al. 2013). Though scant data exist, the lactational transfer of various xenobiotics have been previously described in humans. This includes persistent organic pollutants (Haddad et al. 2015), ellagitannins and their metabolites (Henning et al. 2022), and pharmaceutical drugs and environmental pollutants (Dubbelboer et al. 2023).

### 3.2. The impact of age and diet on the infants’ chemical exposome

To investigate the influence of age and diet on the infants’ chemical exposome, various statistical analyses were applied. For these tests, the raw chromatographic peak areas were utilized following their normalization (SI B). Though this does not accurately take into consideration matrix effects, it is a good compromise for untargeted workflows. In addition, this LC-MS method was previously validated in-house for polyphenols for numerous human bio-fluids (Oesterle et al. 2022, Berger et al. 2024). The first statistical test were principal component analysis (PCA) plots that allowed to investigate the impact of time on the features detected in both matrices (Figures B.6a and b). The breast milk samples showed no clear grouping (Figure B.6a). On the contrary, the infant stool samples showed clear grouping, though there was some overlap between the clusters at months six and twelve (Figure B.6b). The results of both PCAs agree with the hierarchical clustering in the feature heatmaps (Figure 1b and 2b).

As xenobiotics may exhibit synergistic or antagonistic effects, correlations of the features in the breast milk and stool samples were explored. Hence, Spearman rank correlation (p. adj < 0.05) was applied among the 34 identified features in the breast milk (Table A.9, Figure B.5a), and the 68 in the stool (Table A.10, Figure B.5b). In both matrices, the majority of the features positively correlated with one another, especially if they were within the same chemical class. For example, the two lignan microbial metabolites, enterodiol and enterolactone, moderately correlated with each other in both breast milk (ρ = 0.67) and in stool (ρ = 0.52). Of the potentially toxic xenobiotics detected in the stool, the highest correlation was recorded between mono-2-ethylhexyl phthalate and p-nitrophenol (ρ = 0.90). However, no rational for this correlation was apparent. Apart from individual effects, mixture toxicity of these xenobiotics can lead to more severe health consequences especially during early-life (Hamid et al. 2021, Krausová et al. 2023). The only negative correlation observed in the breast milk involved hippuric acid with other phenolic acids, such as benzoic acid (ρ = -0.73), which is most likely due to hippuric acid being a metabolite of benzoic acid (Lees et al. 2013). While in the stool, the only negative correlation was between alternariol monomethyl ether, an *Alternaria* mycotoxin, and genistein-7-sulfate, a metabolite of genistein (ρ = -0.41), thereby indicating potential antagonistic effects between the two. It was previously observed *in vitro* that genistein did have antagonistic effects on the genotoxicity of alternariol, another *Alternaria* mycotoxin (Aichinger et al. 2017). Therefore, further studies are required to explore interactions between xenobiotics *in vivo*. The correlation between the features detected in the breast milk and those in the stool were investigated using the samples from month 1, when the infants were exclusively breastfed. However, due to the limited sample size, no significant correlations were found following Benjamini-Hochberg adjustment. Therefore, future studies with a larger sample size are needed to better characterize the relationship between features detected in the breast milk and corresponding infant stool.

To further investigate changes over time, statistical approaches were applied to the features detected in stool. Firstly, one-way analysis of variance (ANOVA) with false discovery rate post-hoc testing was applied which yielded 325 features that showed significance across all time points (Table A.7). Volcano plots (fold change versus t-test p values) were generated between each time point (Figure B.6c-e), with significant features (fold change > 2, p < 0.05) of each volcano plots listed in Table A.7. A total of 303 features were significant between months one and six, with 151 showing an increase and 153 a decrease. Then from six to twelve months, 118 were significant of which 48 increased and 70 decreased. Finally, from one to twelve months, there were 388 significant features, with 197 increasing and 191 decreasing. ChemRICH plots were then generated with the volcano plot and ANOVA results to depict the chemical classes that were either up- or down-regulated (Figure B.6 f-i). Overall, months one to six and months one to twelve showed up-regulation in chemical classes from plant-origin such as polyphenols, e.g. flavonols or flavones (Figure B.6f-h). Additionally, a down-regulation of chemical classes, such as amino acids and fatty acids, was observed. These results show the alteration in exposure when the infants’ diet changes from milk to complementary foods.

### 3.3. Correlations between features and the developing infant gut microbiome

Spearman rank correlation was applied to preliminarily explore potential correlations between the features that showed significance through ANOVA and the stool microbiome. Several significant correlations between taxa and features in the stool (p. adj < 0.05) were found, and the results (Table A.11) are represented as a heatmap (Figure B.7a) and as a network (Figure B.7b). Specifically, 58 microbe-feature pairs had positive correlations, while 12 microbe-feature pairs showed negative correlations. The correlated features consisted mainly of phytochemicals, for example, the flavone tricetin showed a strong positive correlation with *Blautia* (ρ = 0.69). *Blautia* has been shown to biotransform flavonoids e.g., polymethoxyflavones into demethylated flavones (Kim et al. 2014, Liu et al. 2021). Several flavonoids, including kaempferol and biochanin A, strongly correlated with *Romboutsia* (ρ = 0.70 and 0.68, respectively). An extract containing flavonoids was previously shown to increase the relative abundance of *Romboutsia* in mice (Wang et al. 2022). The betalain vulgaxanthin I showed a strong negative correlation with *Escherichia*-*Shigella* (ρ = -0.70). Extracts containing vulgaxanthin I exhibited antibacterial activities against Gram-negative bacteria, including *Escherichia* (Vulić et al. 2013). Besides phytochemicals, the mycotoxin fumonisin B_1_ showed a strong correlation with *Streptococcus* (ρ = 0.75). Previously, it was observed that *Streptococcus* can bind to fumonisin B_1_ (Niderkorn et al. 2006). While the causal link between the features derived from xenobiotics and members of the infant gut microbiome remains to be elucidated, the results highlight the need to explore xenobiotic-microbiome interactions in less complex *in vitro* models and more detail.

### 3.4. Limitations

There are several limitations to consider in the applied analytical workflow and the overall study design. Many of the analytes investigated, especially polyphenols, have a wide range of isomers, making annotation of the features a complex task. This is reflected in the suspect screening results where many features (Table A.6 and A.7) have the same annotation though they were distinct analytes. Additionally, DIA MS^2^ spectra typically have more noise than DDA MS^2^ spectra (Guo and Huan 2020), further complicating annotation, especially when using *in silico* fragmentation. The uncertainty in feature annotation also impacts the biological relevance of the statistical and correlation analyses. Furthermore, the correlation and statistical results must be interpreted with caution due to the limited number of participants in the study (n = 11). As the sample size was low, all time points were included in the correlations between the infant stool and gut microbiome. Therefore, the correlations may reflect indirect drivers such as the dynamic development of the microbiome in early life or changes in diet with age.

## 4. Conclusion

Exposure to natural and anthropogenic chemicals during early life is known to have a considerable impact on the development and health of humans. This study provides a preliminary overview of the presence of potentially beneficial as well as potentially toxic xenobiotics in breast milk and stool from Nigerian mother-infant pairs. Several xenobiotics detected in the breast milk were also present in the corresponding stool samples, although the stool samples contained, as expected, a higher number of diverse xenobiotics. Infant exposure to xenobiotics significantly increased with the introduction of complementary foods. Moreover, correlations were observed between xenobiotics and certain members of the gut microbiome of the infants. However, the toxicological relevance of these results needs to be further explored in larger cohorts and validated in *in vitro* models. Data is sparse on longitudinal metabolomic/exposomic profiles of healthy Nigerian mother-infant pairs. Thereby, despite the limited sample size, the data herein gives an important snapshot of the chemical exposome in biofluids of a population that has been considered to be at relatively high exposure levels to many beneficial and adverse xenobiotics. The next steps should be the application of such workflows in larger cohorts and different populations, especially in long-term studies, to better characterize the influence that exposure to various chemicals has on health and microbiome development.

## Acknowledgment

The authors would like to thank the mothers and their infants for providing the samples. They would also like to thank and acknowledge all the members of their working groups, the Warth and Rompel labs, for their help, support, and feedback. The authors would like to express their gratitude to the Mass Spectrometry Center of the Faculty of Chemistry at the University of Vienna for technical support during the measurements, and to the Joint Microbiome Facility of the University of Vienna and the Medical University of Vienna. This work was supported by the University of Vienna through the Exposome Austria Research Infrastructure, the Austrian Federal Ministry of Education, Science and Research (project DigiOmics4AT, B.W.), the Austrian Federal Ministry for Climate Protection, Environment, Energy, Mobility, Innovation and technology (BMK), and the Austrian Science Fund (FWF) [10.55776/P32326 and 10.55776/P32932] (to A.R.).

For open access purposes, the authors have applied a CC BY public copyright license to any author-accepted manuscript version arising from this submission.

## Data availability

LC-MS raw data files have been submitted to the MetaboLights data repository (MTBLS8792). The 16S rRNA gene amplicon data is available on the BioProject accession number PRJNA1013123.

## Appendix

Supplementary file A (Excel) contains all of the tables mentioned in the text, e.g. the suspect screening results for breast milk and stool.

Supplementary file B (PDF) contains further details on the materials and methods, and additional figures, including boxplots of the semi-quantification results and heatmaps from the correlation analysis.

## CRediT author contributions

**Ian Oesterle:** Conceptualization, Methodology, Software, Formal analysis, Investigation, Writing - Original Draft, Writing - Review & Editing, Visualization. **Kolawole I. Ayeni:** Conceptualization, Software, Formal analysis, Writing - Review & Editing. **Chibundu N. Ezekiel:** Conceptualization, Writing - Review & Editing, Supervision, Funding acquisition, Resources. **David Berry:** Writing - Review & Editing, Supervision. **Annette Rompel:** Writing - Review & Editing, Supervision, Funding acquisition, Resources. **Benedikt Warth:** Conceptualization, Writing - Original Draft, Writing - Review & Editing, Supervision, Funding acquisition, Resources.

