## Supplementary File B for "Insights into the early-life chemical exposome of Nigerian infants and potential correlations with the developing gut microbiome"

### Table of Contents

1. **Sample preparation**…………………………………………………………………...Page 2
2. **Data processing of the acquired LC-HRMS data**……..………...…………...……..…..…Page 2
3. **Quality control of the LC-HRMS measurements**…………………………………...……Page 2
4. **Quantification of Level 1 identified features**……………………………………...…Page 3
5. **Statistical analysis**……………………………...……………………………...…Page 3
6. **Figure B.1.** Extracted Ion Chromatograms of Daidzein and Daidzein-7-ß-d-glucuronide.…Page 4
7. **Figure B.2.** Heatmap with Ward clustering of all the features detected in the breast milk and stool samples. ……………………..……………………...……………………………...…Page 5
8. **Figure B.3.** Box plots of the features identified and semi-quantified with reference standards in the breast milk samples at the different time points separated by their chemical class. ..……Page 6
9. **Figure B.4.** Box plots of the features identified and semi-quantified with reference standards in the stool samples at the different time points separated by their chemical class. ..…………Page 13
10. **Figure B.5.** Spearman rank correlation of identified features. ..…………………………....Page 23
11. **Figure B.6.** Statistical analysis of the features detected in the stool samples.......…...........Page 24
12. **Figure B.7.** Correlations between features in the stool sample and the gut microbiome..…Page 25
13. **References**…………………………………………………………………….…Page 26

### 1. Sample preparation

The infant stool samples were prepared following a slightly modified method from Krausová et al. (2022). In brief, approximately 80 mg of the wet infant stool was weighed in a micro-reaction tube and dried in a vacuum concentrator (Labconco). Water ($H_2O$) was then added to the samples at 40 µL per 20 mg of dried stool, followed by 160 µL per 20 mg of dried stool of acetonitrile (ACN): methanol (MeOH) (1:1) + 1% v/v formic acid (FA). The samples were then vortexed and ultrasonicated on ice for 15 min, and subsequently placed at -20°C overnight to allow for protein precipitation. After, they were centrifuged for 10 min at 4°C and 18,000 x g. The supernatant was transferred to a new micro-reaction tube and diluted 10 times with ACN:$H_2O$ (1:1) + 1% v/v FA, and finally the samples were passed through a PTFE filter into an amber LC vial.

### 2. Data processing of the acquired LC-HRMS data

The acquired raw data files were first converted to ABF file format with Reifycs Analysis Base File Converter (v2011-2020), before they were further processed in MS-Dial (v4.9.221218) (Tsugawa et al. 2015). MS-Dial was used for feature pre-processing, e.g., building extracted ion chromatograms, and for feature annotation with a spectral library created by MS-Dial that combines various databases such as GNPS and MassBank (Tsugawa et al. 2015). The parameters used in MS-Dial for the two biological matrices are listed in Table A.4. Features were further annotated with *in silico* fragmentation using MS-Finder (v3.52) (Tsugawa et al. 2016). Prior to feature annotation, the feature lists were filtered by $MS^1$ matching with two online databases: PhytoHub which contains 2268 phytochemicals, mainly polyphenols (Giacomoni et al. 2017), and Exposome-Explorer, which contains 1262 chemicals known to be biomarkers for exposure to environmental and lifestyle factors such as diet, pollutants, or contaminants (Neveu et al. 2020). R (v4.3.1) (R Core Team 2023) was used for $MS^1$ matching and feature clean-up with the process blank and pooled quality control (QC) samples.

The identification levels previously defined by Schymanski et al. (2014) were applied. In brief, features identified with authentic reference standards were labeled as Level 1 and features annotated with spectral libraries as Level 2a. Level 3 was then split in a similar manner as described in Oesterle et al. (2023), with features annotated by in silico fragmentation were labeled as Level 3a, and features putatively annotated by their $MS^1$ with the two databases as Level 3b. The chemical classes of the features were also determined using the ChemRICH MeSH prediction tool (Barupal and Fiehn 2017), the classes listed in MS-Finder and MS-Dial, and the classes listed in the entries of the two online databases (PhytoHub and Exposome-Explorer).

### 3. Quality control of the LC-HRMS measurements

QC samples were prepared for each biological matrix by pooling aliquots of the respective samples. The stool QC was prepared by pooling aliquots of each infants' wet stool in respective quantities (Table A.2). The breast milk QC was prepared by combining 10 µL of each sample. The QC samples were processed following the same procedure as the experimental samples. For each biological matrix, the respective QC was used to condition the LC column prior to the acquisition of the experimental samples. Moreover, after column conditioning, three technical replicates of the QCs were measured. The QCs were routinely analyzed after every five experimental samples to continuously check the reliability of the instrument as well as to correct for any signal drifts that may occur within the acquisition batch. In addition, for each QC, a five-point serial dilution series with a constant dilution factor of four was prepared with the respective dilution solvent of the biological matrix's sample preparation procedure. For each biological matrix, along with the samples, a process blank was prepared by leaving the micro-reaction tube empty rather than taking an aliquot of the biological matrix. For the stool protocol, the process blank was assumed to contain 20 mg of dry weight, therefore 40 μL of $H_2O$ and 160 μL of ACN:MeOH (1:1) + 1% v/v FA were added. Both process blanks were measured in triplicates.

Features with an average chromatographic peak area in the process blank measurements greater than one-third of the average in the samples were removed (Kirwan et al. 2014). In addition, features were removed unless they were detected in at least two of the triplicate measurements of the QCs and their peak area in the QC triplicate measurements had a relative standard deviation <30% (Dudzik et al. 2018). Features with an average signal-to-noise value of less than three, as calculated by MS-Dial, were also removed. Finally, the QC dilution series was developed to assess the reliability of extracted features, meaning a feature's signal in the QCs should decrease as the QC is diluted. Therefore, Spearman rank correlation was applied to the QC dilution series for assessing the association between a feature´s signal and the overall dilution factor, whereby features with a correlation value less than 0 were removed.

### 4. Quantification of Level 1 identified features

The identified features (Level 1) were semi-quantified with calibration curves created from the reference standards (Table A.5). To create the calibrations curves, the pooled QC for each respective matrix was spiked with the reference standards at different concentrations. The signal suppression and enhancement effects were thus approximated by making matrix-matched calibration curves. However, as the pooled QC was used for spiking, the Level 1 identified features already had a chromatographic peak present. Therefore, standard addition was applied for quantification.

### 5. Statistical analysis

In MetaboAnalyst (v5.0) (Pang et al. 2022), the features in each matrix were both normalized by median and applied $\log_{10}$ transformation. Then, principal component analysis (PCA) of the features were created to investigate clustering of the samples in each matrix respectively. For the stool samples, one-way analysis of variance (ANOVA) was then applied to determine significant features over time, while volcano plots (fold change versus T-tests) were used to determine fold changes between the time points. Heatmaps with Ward hierarchical clustering of the annotated features were created to compare the distribution in each matrix.

Boxplots of the quantification results were made with the *ggplot2* (v3.4.3) (Wickham 2016) and *cowplot* (v1.1.1) (Wilke 2020) packages. Spearman rank correlation between the features in both matrices was calculated using *stats* (v3.6.2) package in R (v4.3.1) (R Core Team 2023). The *pheatmap* package (v1.0.12) (Kolde 2018) was applied to generate heatmaps of the correlation results. Benjamini-Hochberg was applied for multiple testing (Benjamini and Hochberg 1995).

Microbiome data was loaded into R (v4.3.1) and filtered using the *ampvis2* package (v2.8.3) (Andersen et al. 2018). Spearman rank correlations with Benjamini-Hochberg correction between features and the microbiome was done using the *stats* (v3.6.2) package and visualized as heatmaps with the *pheatmap* (v1.0.12) package. To further visualize the correlations between the features and the microbiome, a network was created in Cytoscape (v3.9.1) (Shannon et al. 2003).

**6. Figure B.1.** Extracted Ion Chromatograms of Daidzein and Daidzein-7-ß-d-glucuronide.

Extracted ion chromatograms of daidzein (m/z 253.050, retention time: 7.7 min, Level 1) (**A**) and daidzein-7-ß-d-glucuronide (m/z 429.079, retention time: 5.6 min, Level 1) (**B**) in all the infants at the three distinct time points. In addition, the process blank, pooled QC, and pooled QC spiked with the authentic reference standards (14 ng/mL for daidzein and 1.1 ng/mL for daidzein-7-ß-D-glucuronide) are depicted. These chromatograms illustrate that xenobiotics of plant origin significantly increase in concentrations as complementary foods are introduced in the infants' diets

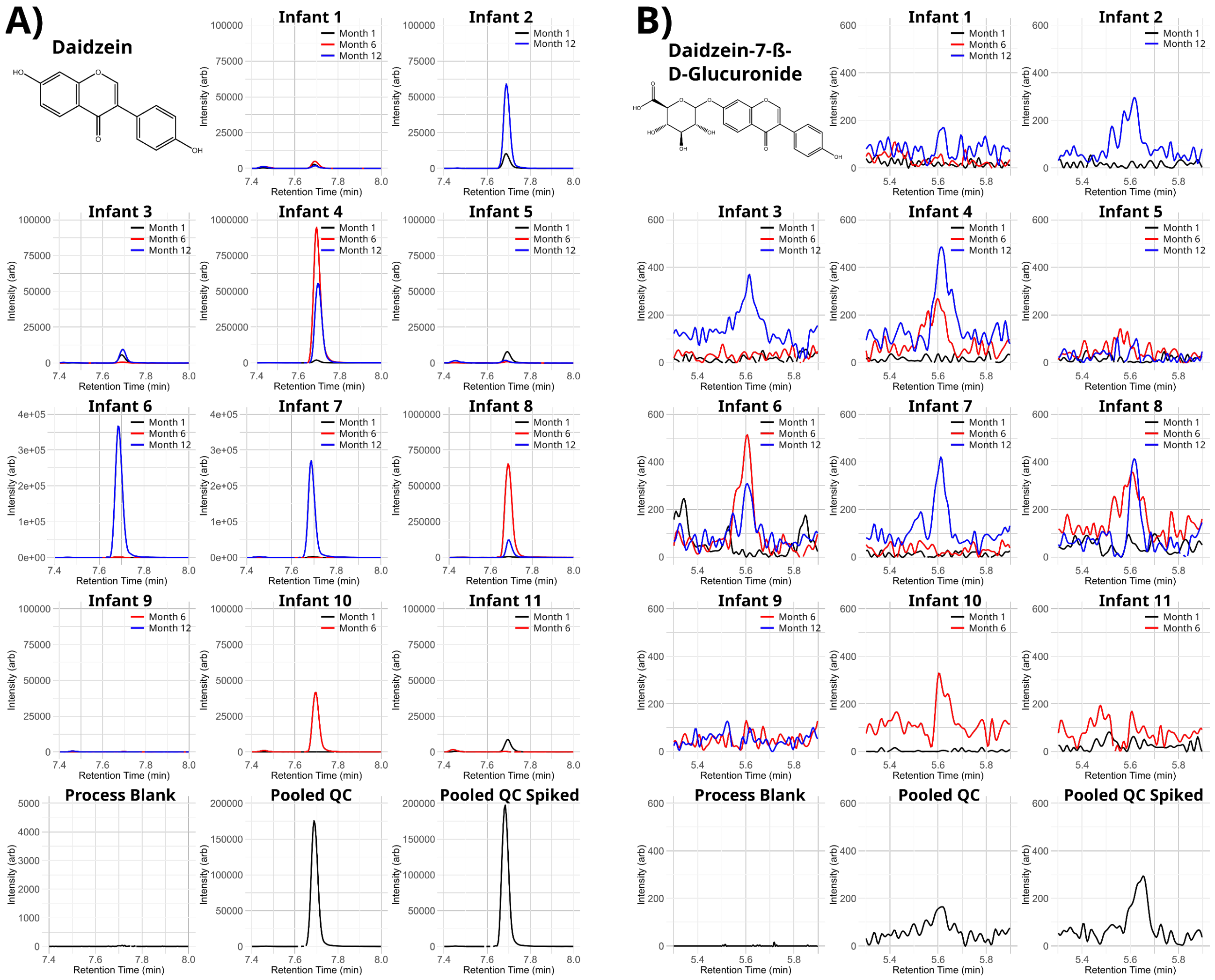

**7. Figure B.2.** Heatmap with Ward clustering of all the features detected in the breast milk and stool samples.

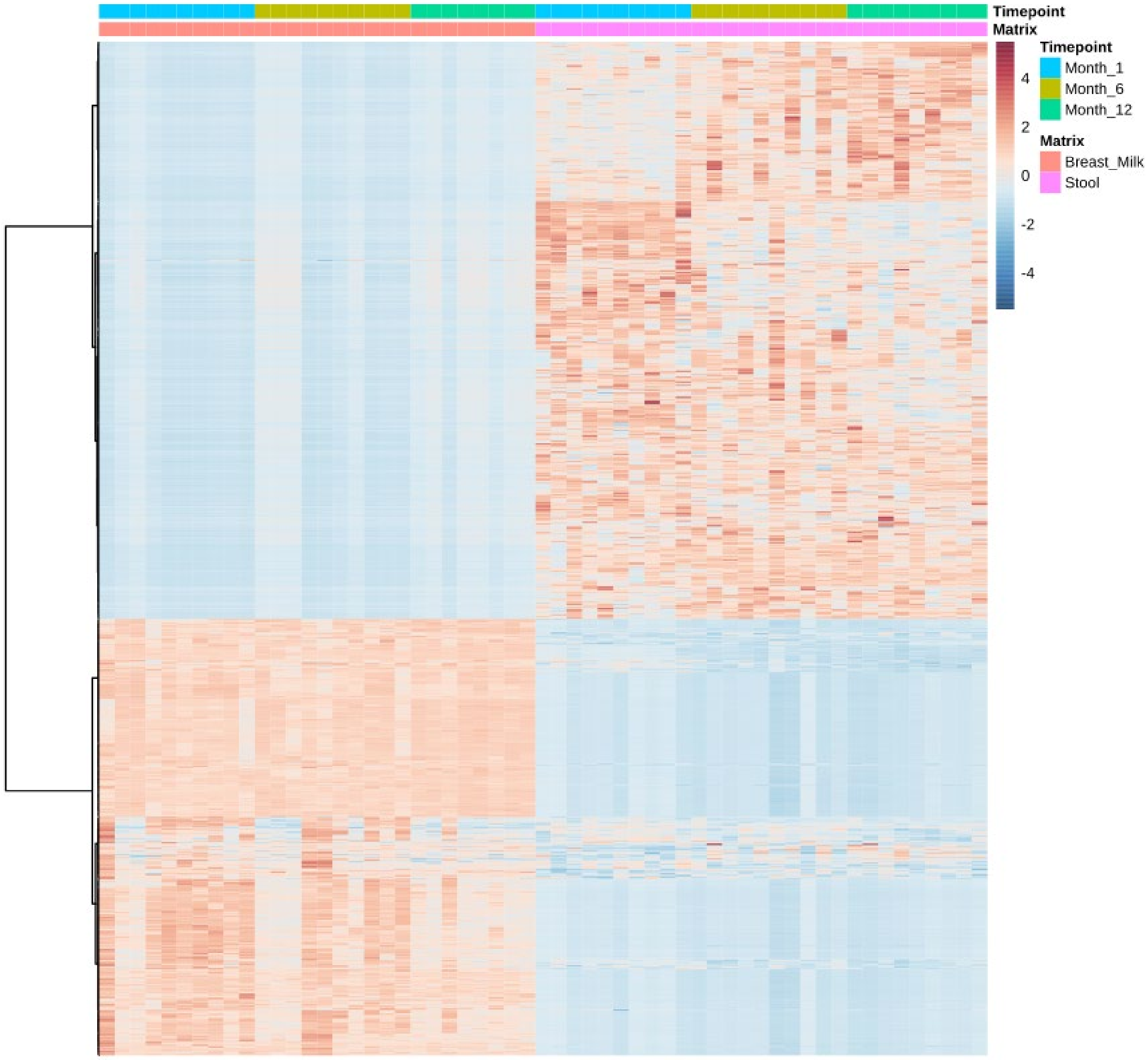

**8. Figure B.3.** Box plots of the features identified and semi-quantified with reference standards in the breast milk samples at the different time points separated by their chemical class.

**Chalcones**

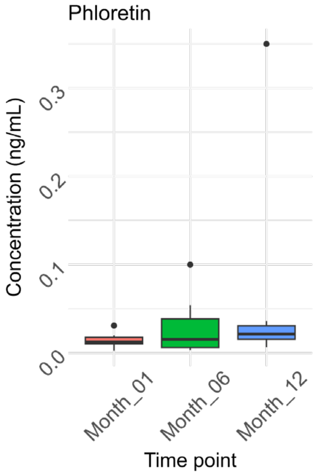


**Flavanones**

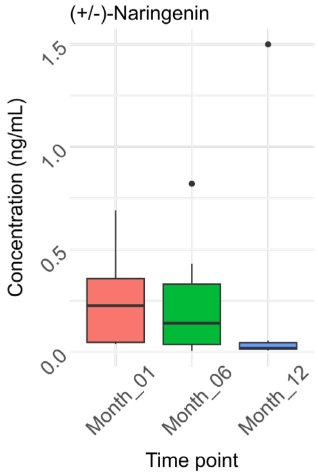


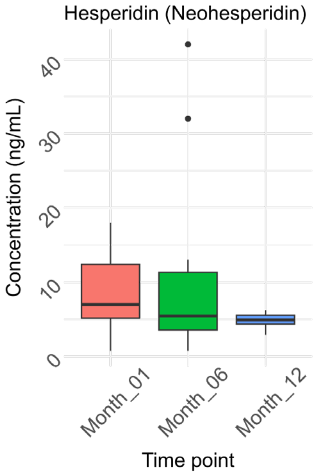


**Flavanols**

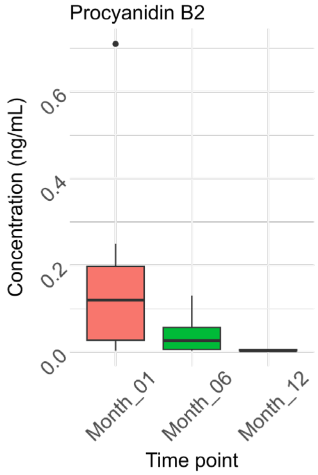

### Flavones

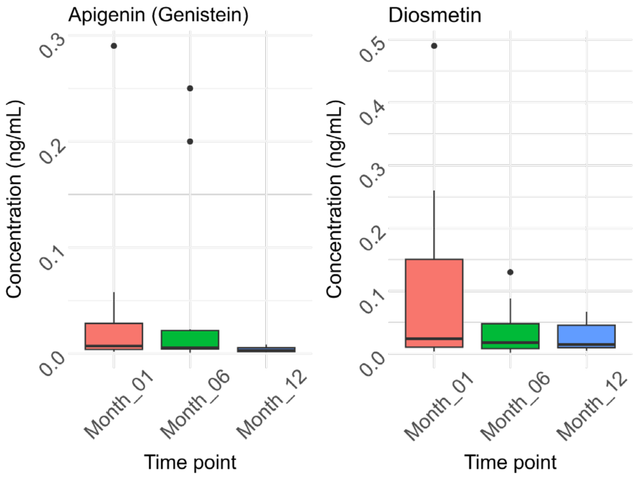


### Isoflavones

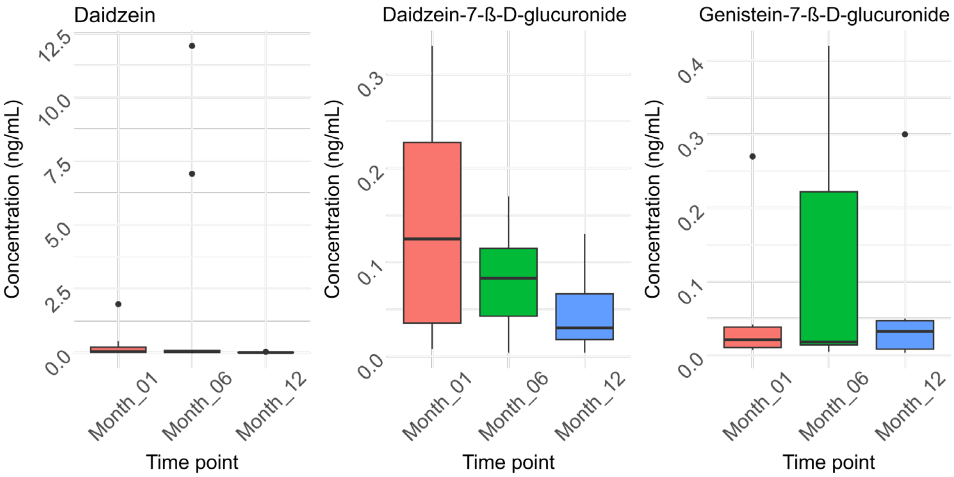


### Lignans

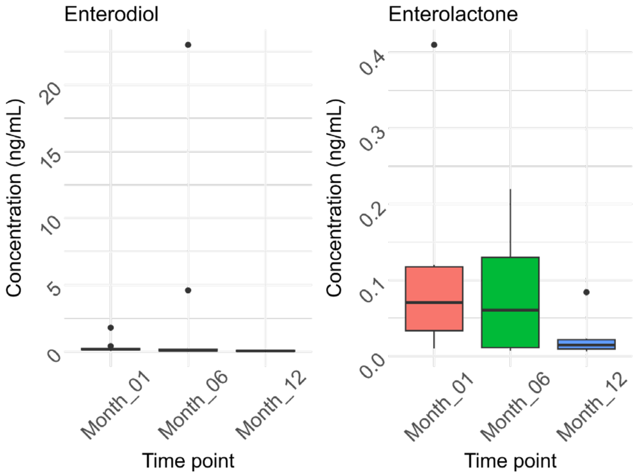

### Benzoic and hippuric acids

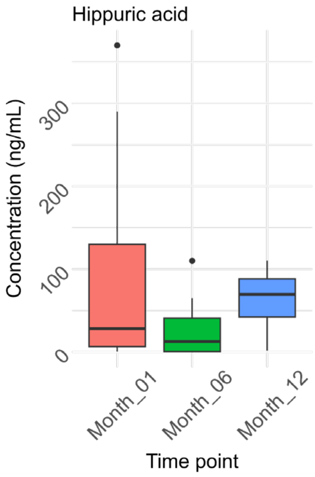


### Hydroxybenzoic acids

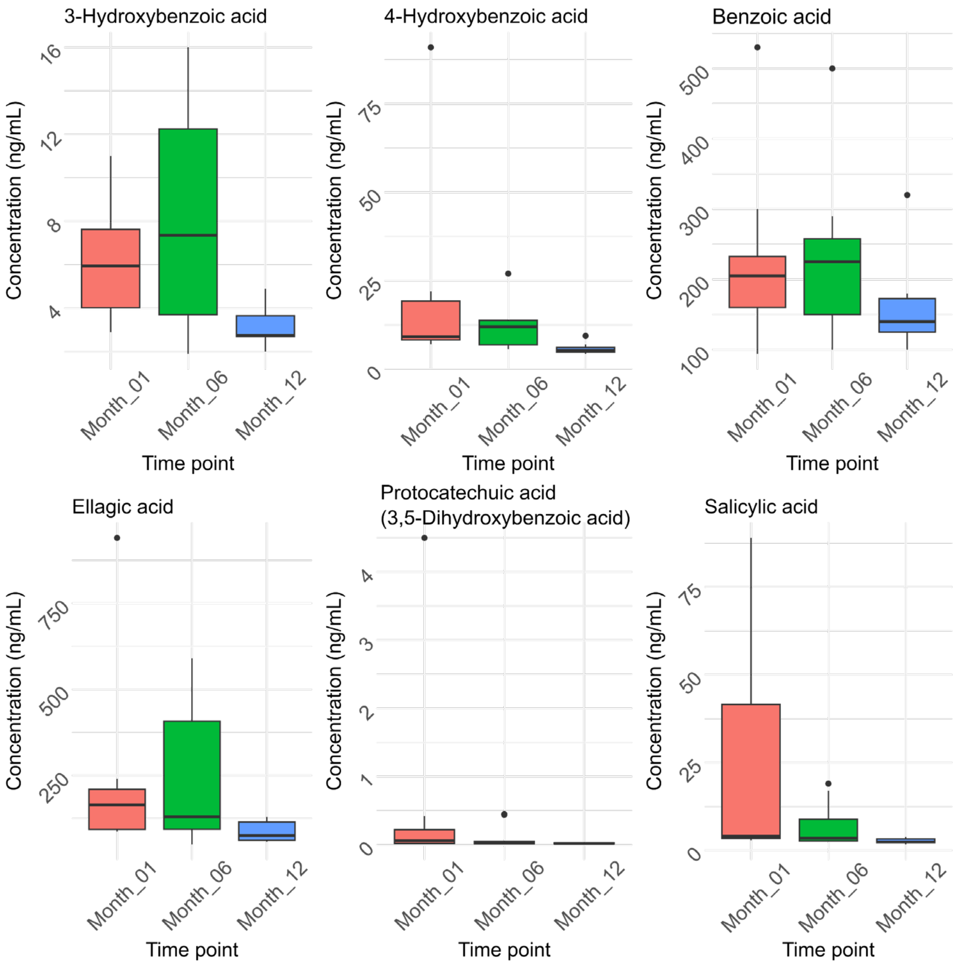

### Hydroxycinnamic acids

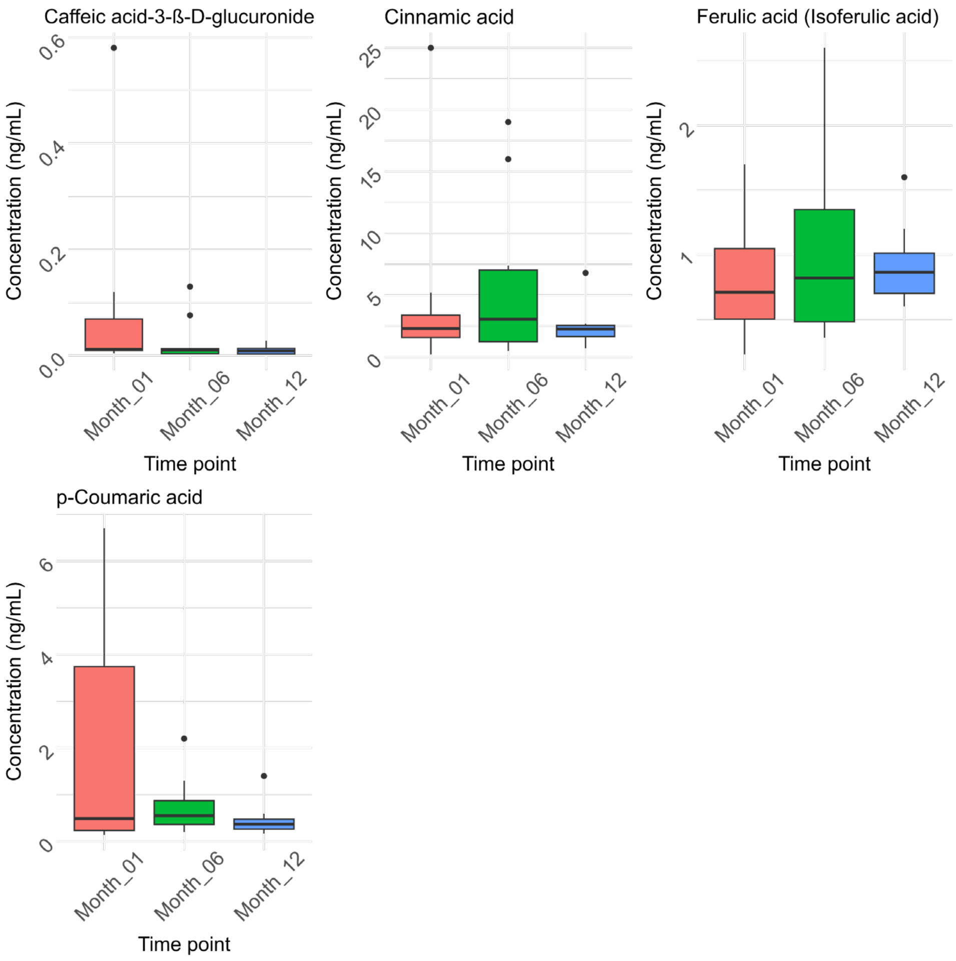


### Phenolic acids

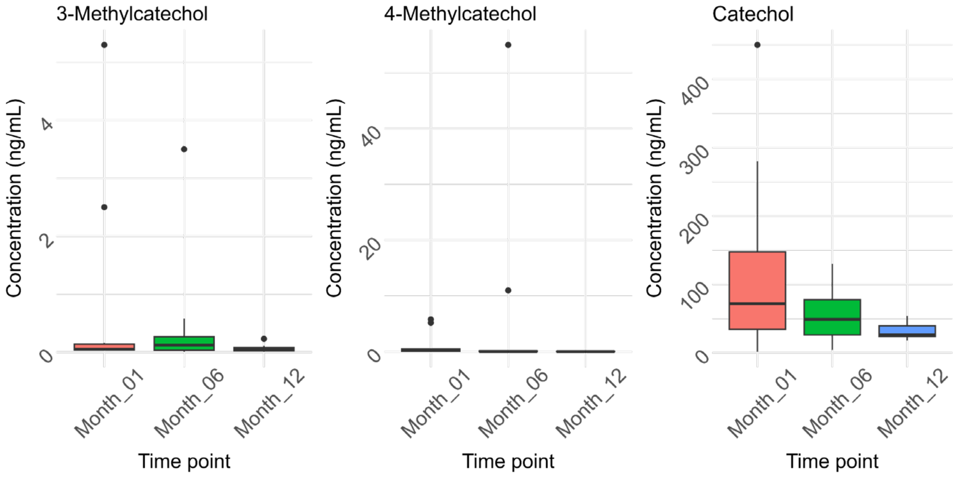

### Phenylacetic acids

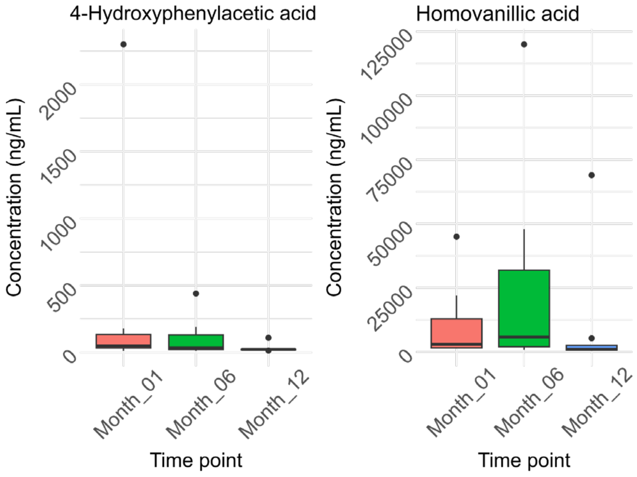


### Phenylpropanoic acids

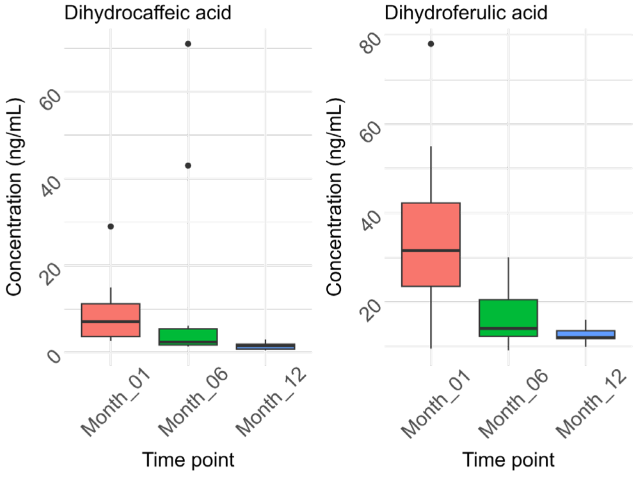


### Stilbenes

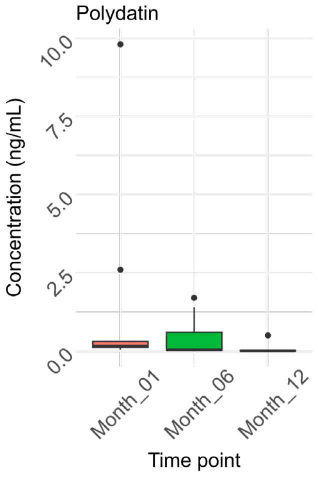

### Antibiotics

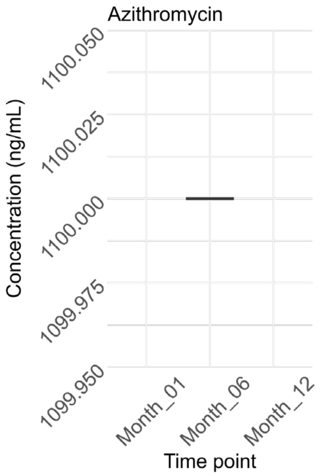


### Estrogens

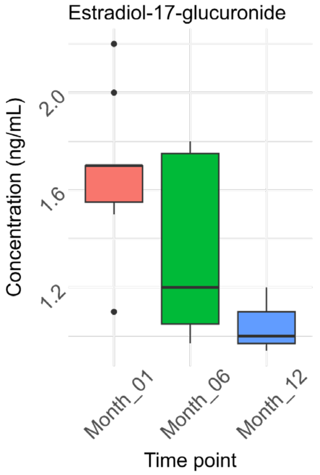


### Mycotoxins

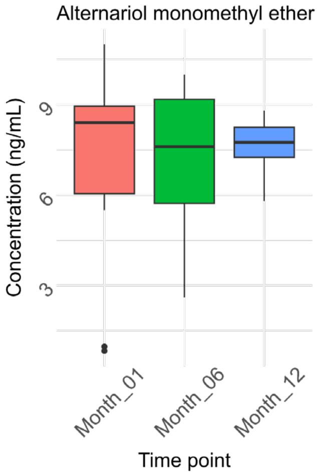

**Personal care product / Pharmaceutical ingredients**

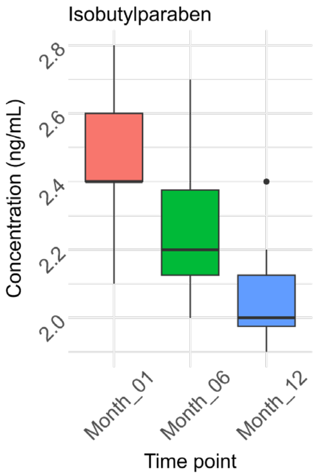

**9. Figure B.4.** Box plots of the features identified and semi-quantified with reference standards in the stool samples at the different time points separated by their chemical class.

**Catechins**

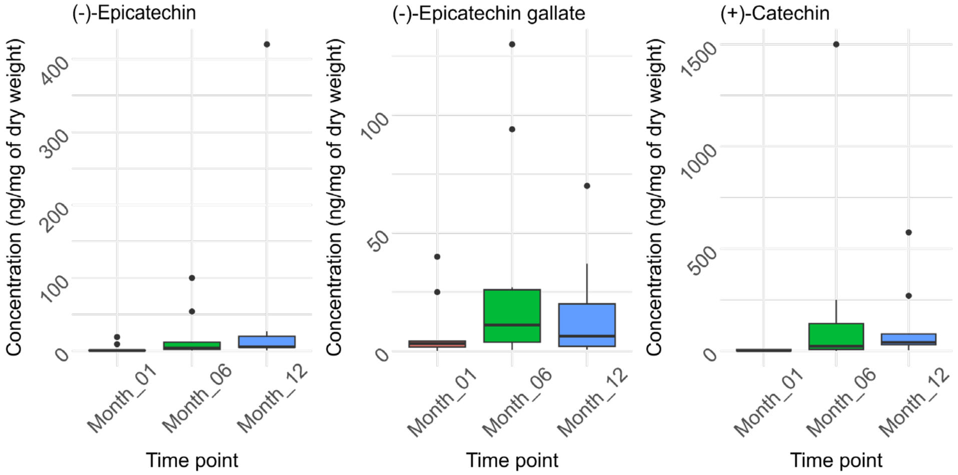


**Chalcones**

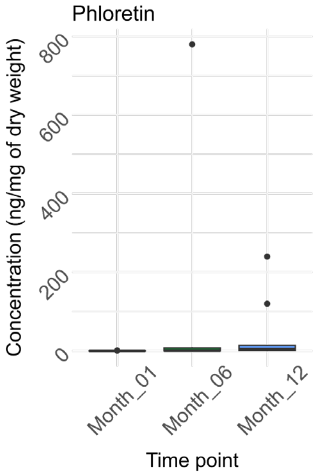


**Flavanones**

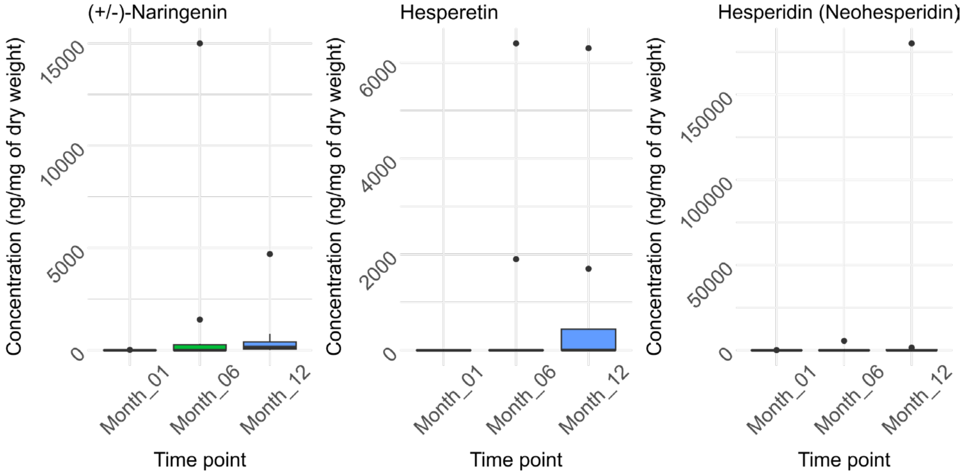

### Flavanols

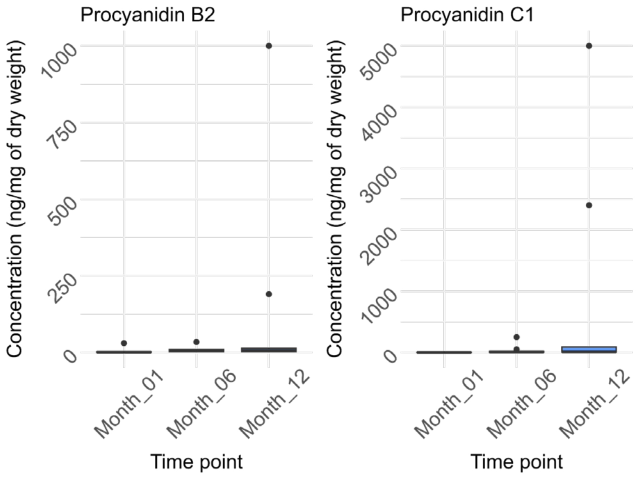


### Flavones

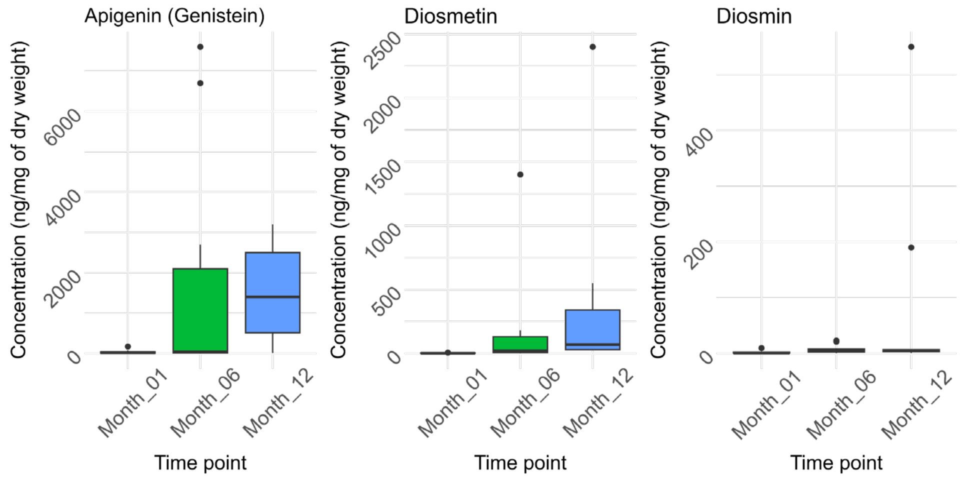


### Lignans

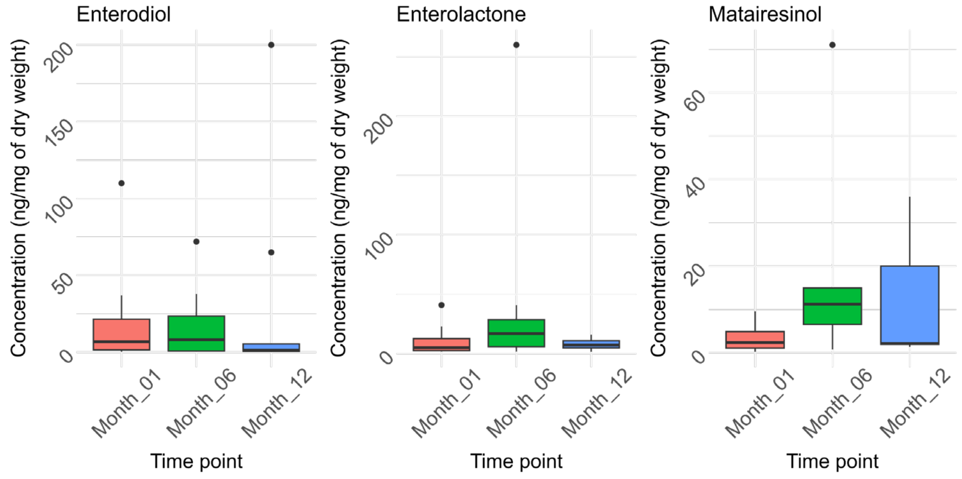

### Flavonols

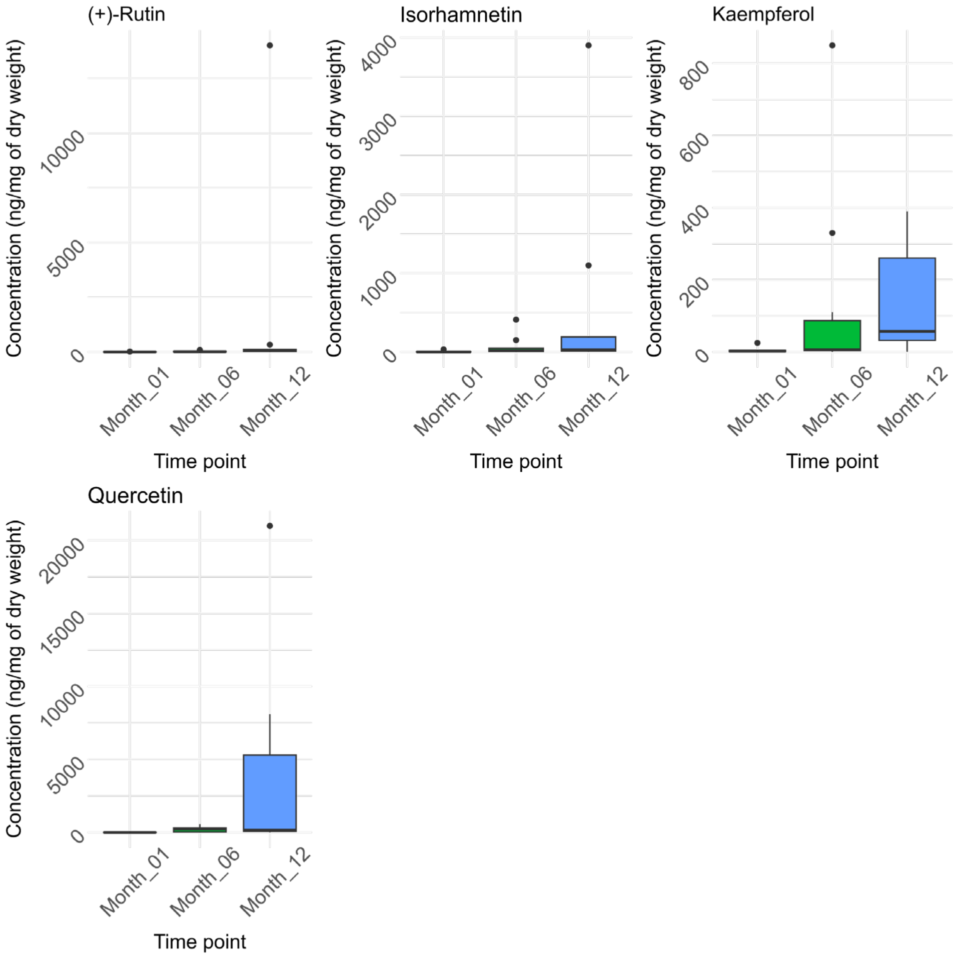


### Benzoic and hippuric acids

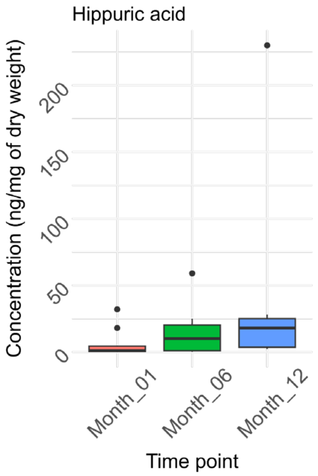

### Isoflavones

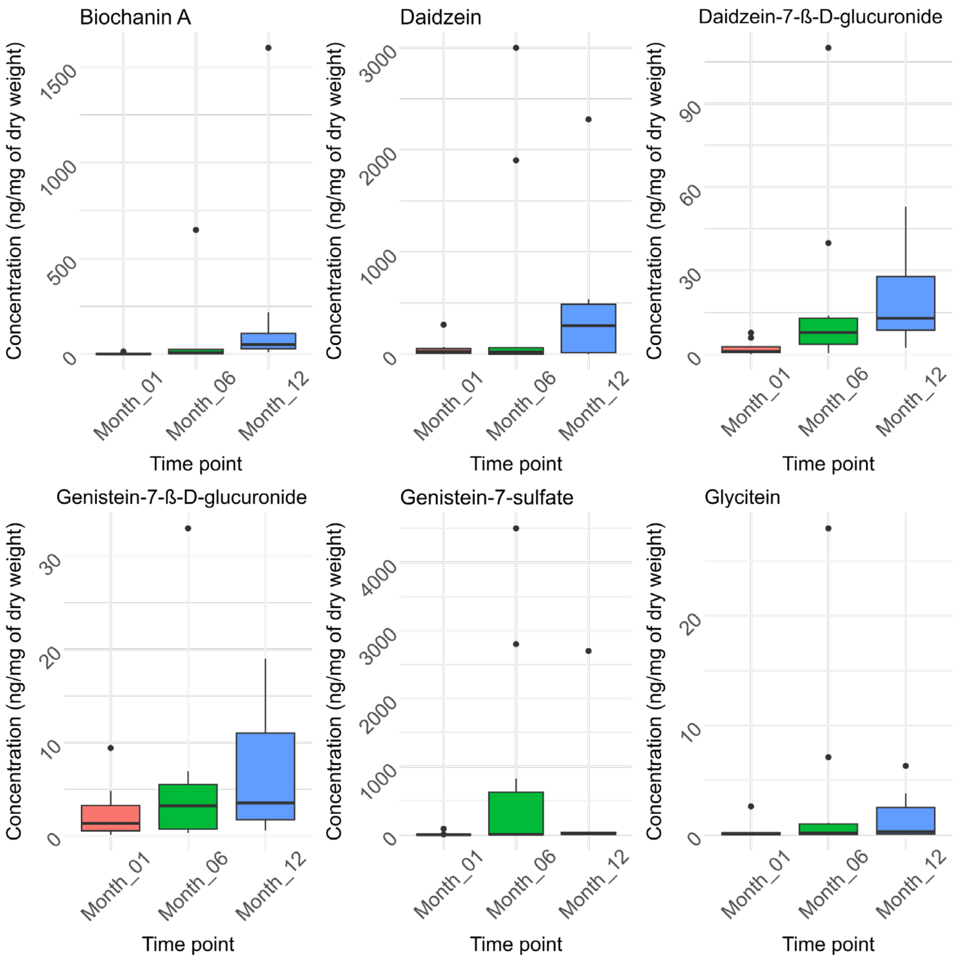


### Phenolic acids

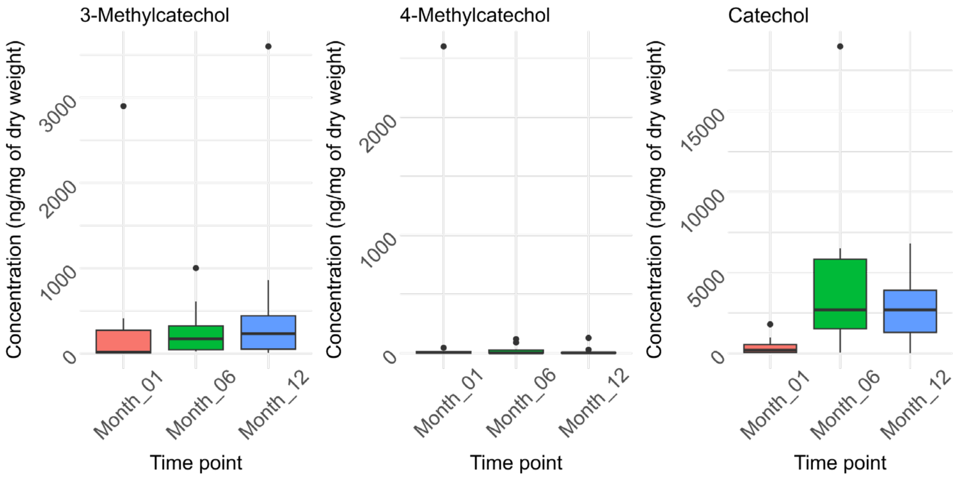

### Hydroxybenzoic acids

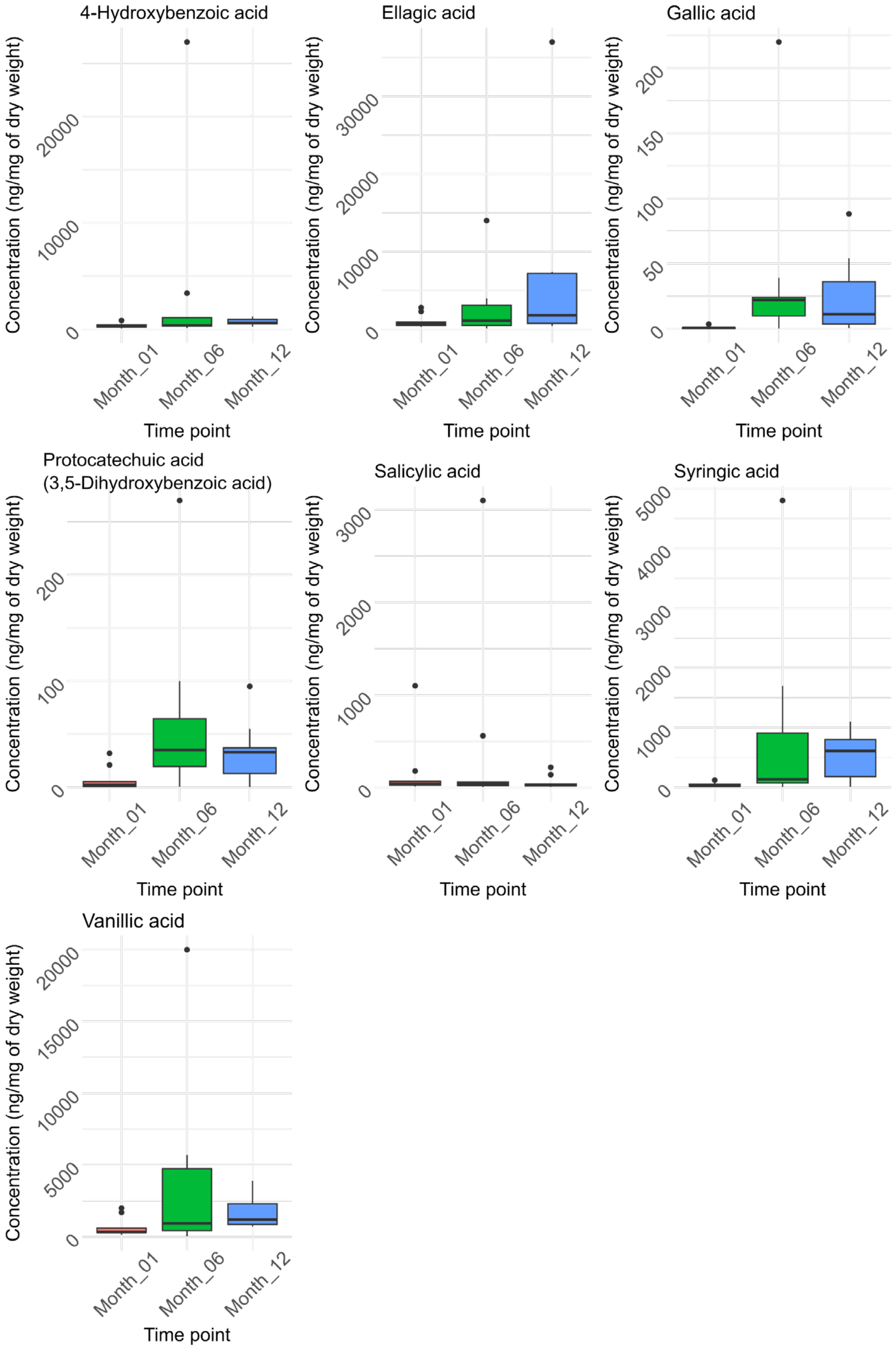

### Hydroxycinnamic acids

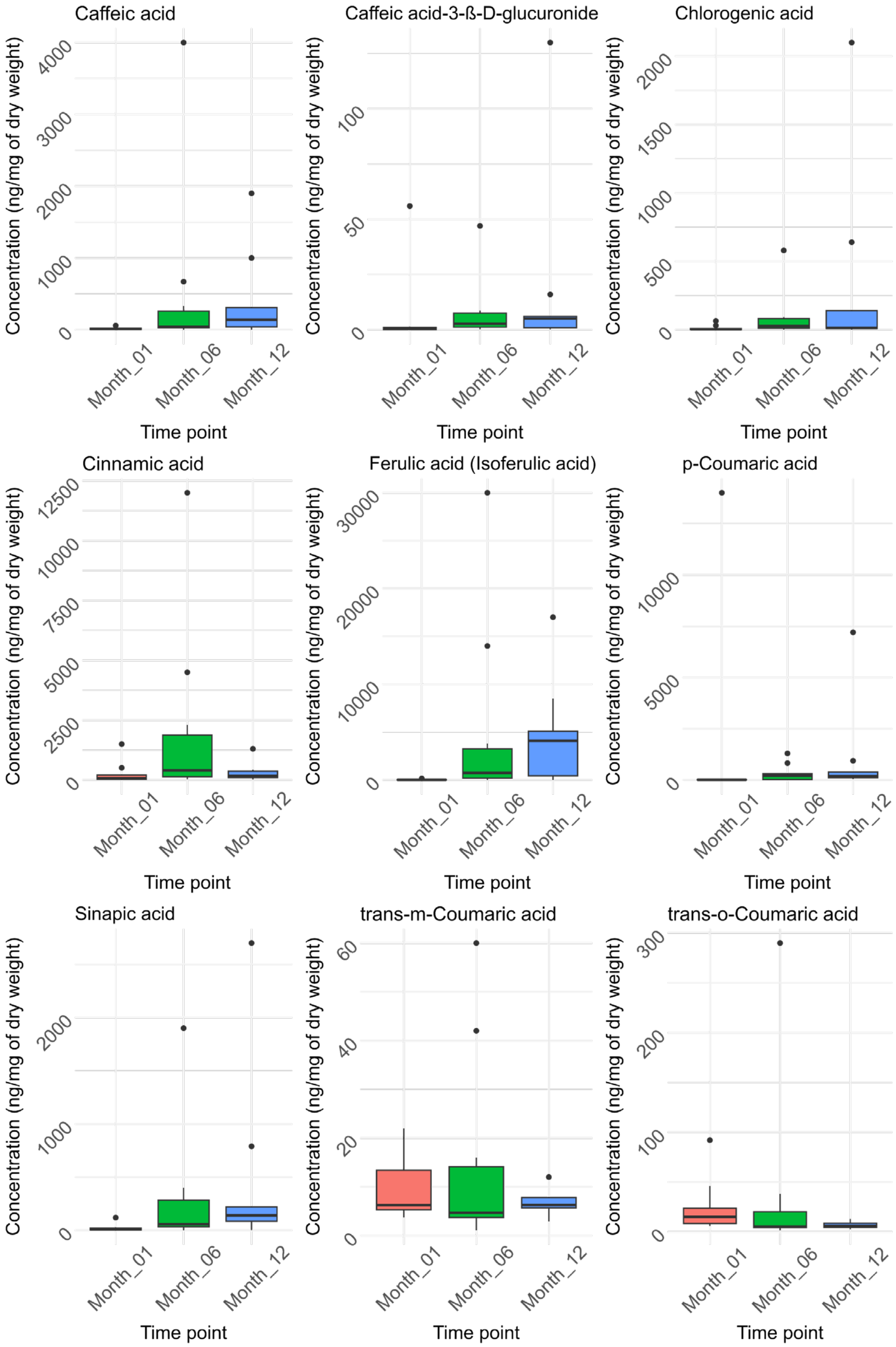

### Phenylacetic acids

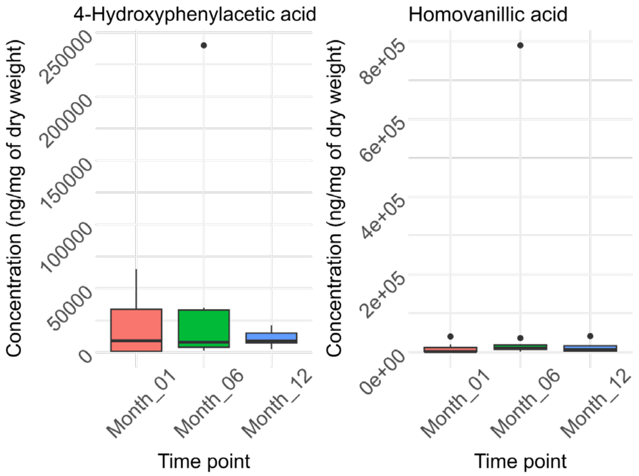


### Phenylpropanoic acids

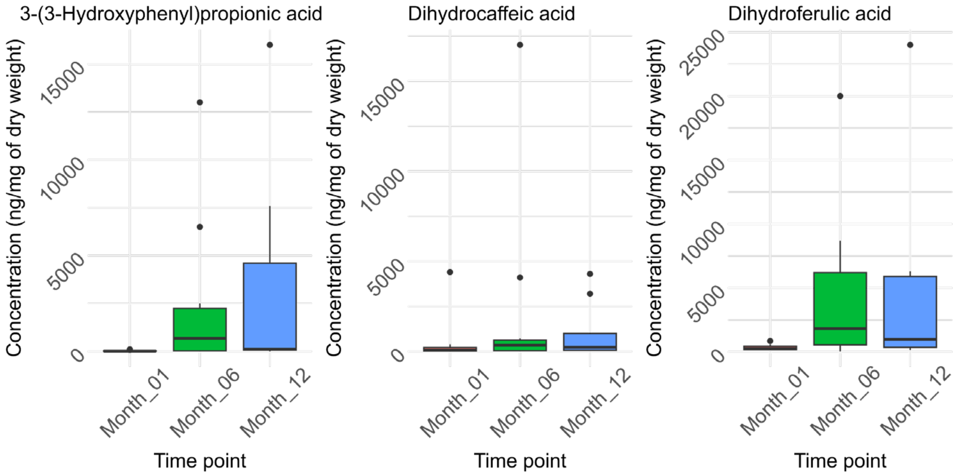


### Stilbenes

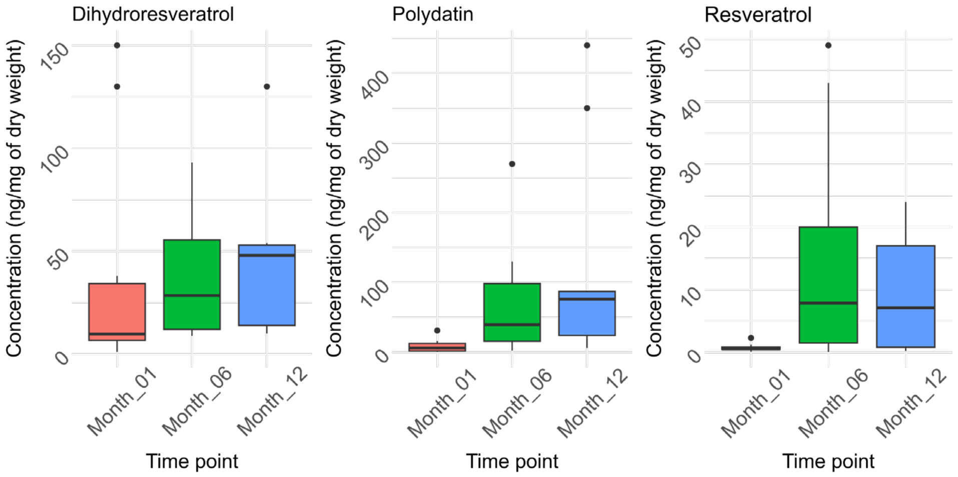

### Urolithins

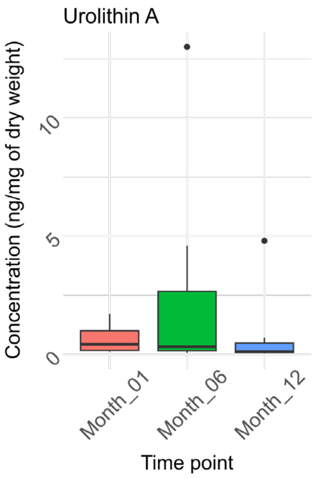


### Mycotoxins

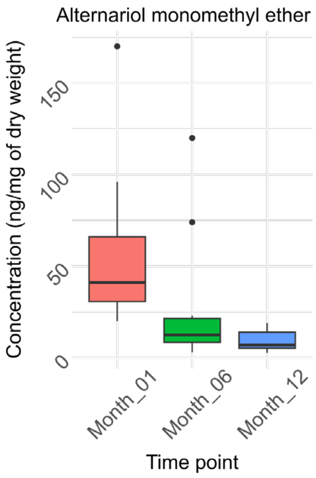


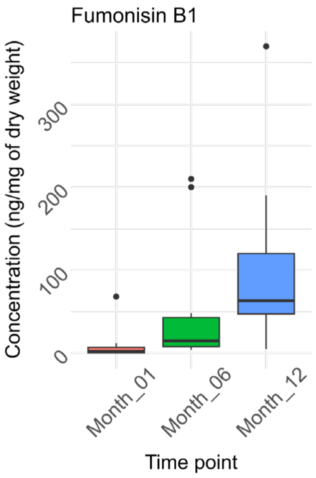


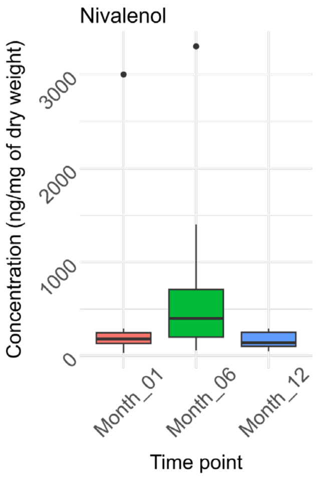


### Pesticides

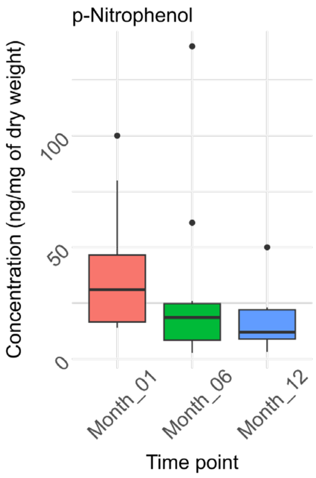

### Antibiotics

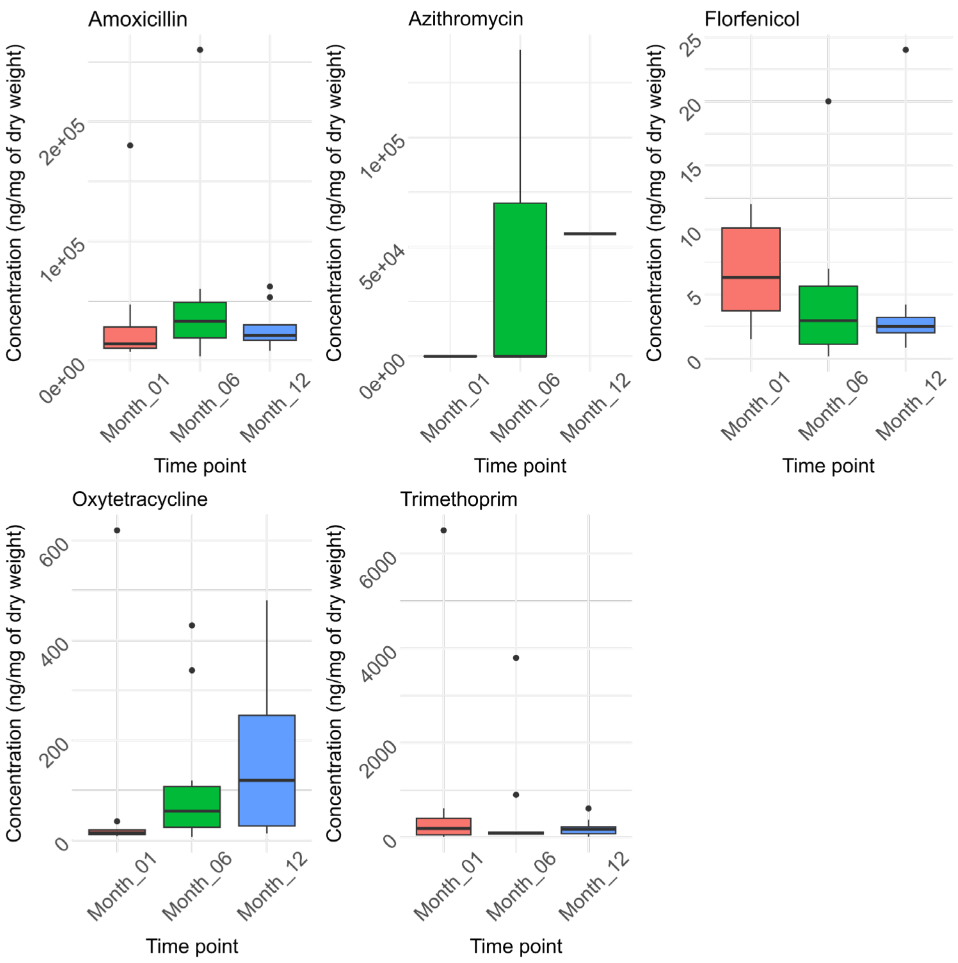


### Personal care product / Pharmaceutical ingredient

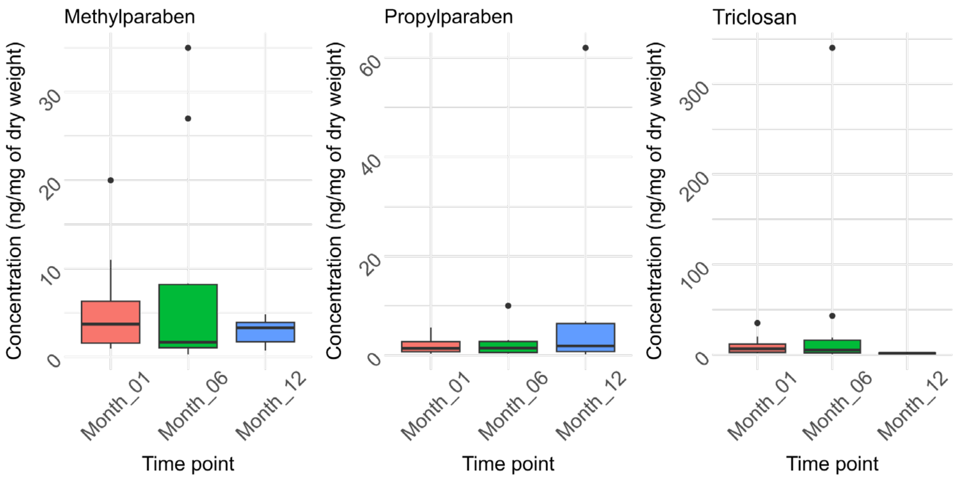

### Perfluorinated alkylated substances

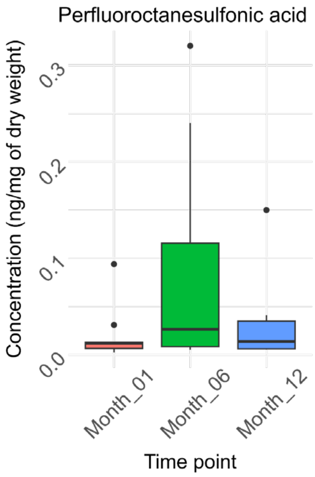


### Plasticizer / Plastic component

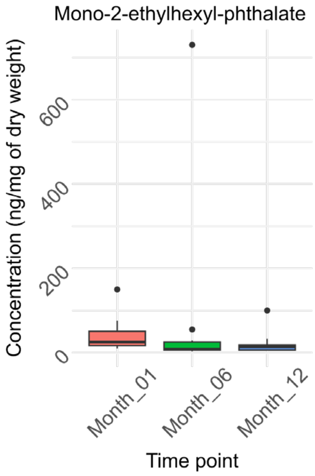

**10. Figure B.5.** Spearman rank correlation of identified features.

**A)** Spearman rank correlation (p. adj < 0.05) heatmap between the 34 identified features (Level 1) in the breast milk samples. **B)** Spearman rank correlation (p. adj < 0.05) heatmap between the 67 identified features (Level 1) in the stool samples.

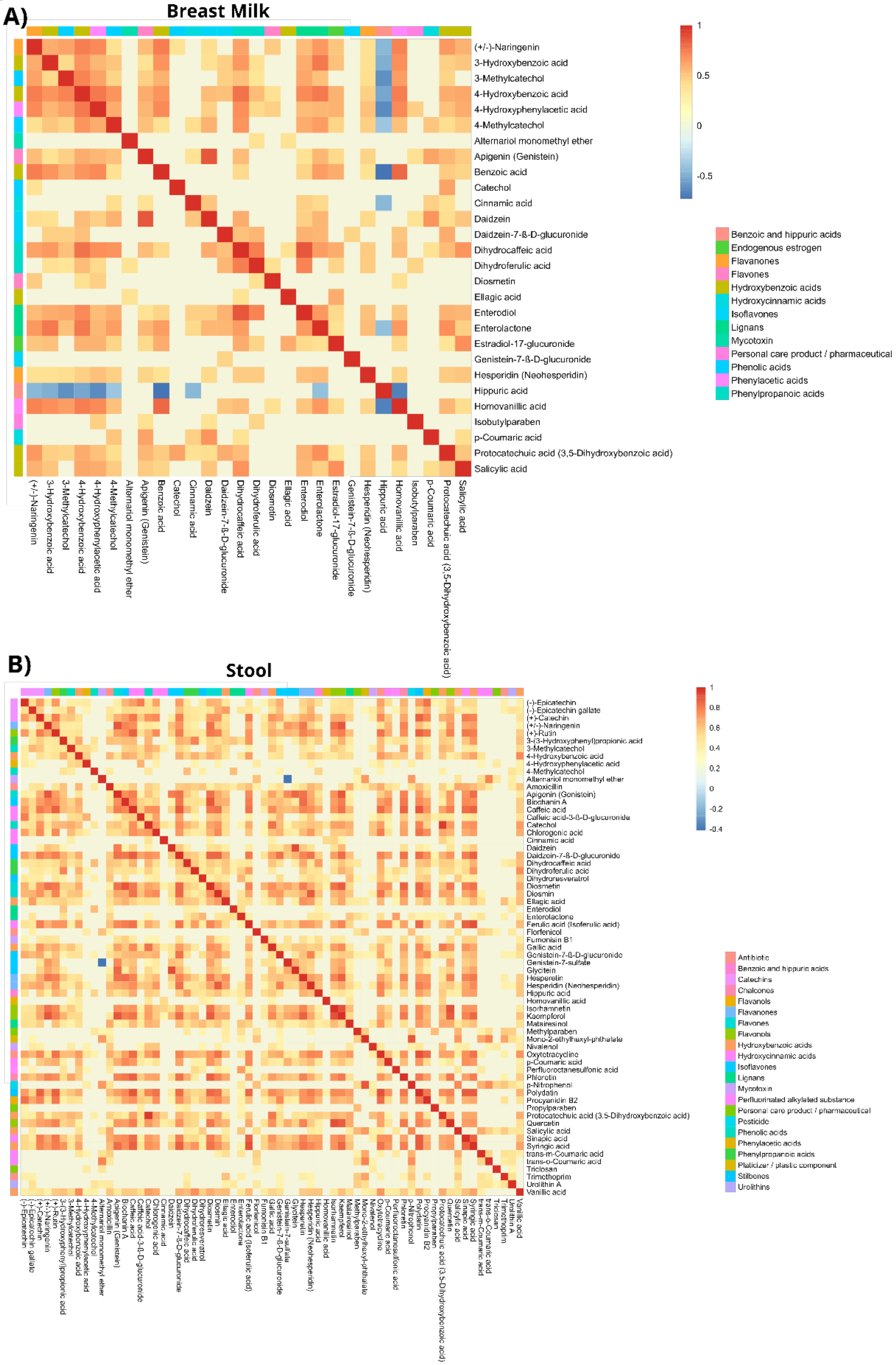

**11. Figure B.6.** Statistical analysis of the features detected in the stool samples.

**A)** Principal component analysis (PCA) plot of the breast milk samples demonstrates that the milk composition did not cluster, indicating no major overall changes of the breast milk composition over time. This makes the study design suitable to explore the impact of solid food intake introduced during early life. **B)** PCA plot of the infant stool samples shows grouping by age with a clear separation of the first month after delivery. **C)** Volcano plot ($p < 0.05$, fold change > 2) of the features between the samples from infants at one month old and six months old. **D)** Volcano plot ($p < 0.05$, fold change > 2) of the features between the samples from infants at one month old and twelve months old. **E)** Volcano plot ($p < 0.05$, fold change > 2) of the features between the samples from infants at six months old and twelve months old. **F)** Chemical enrichment (ChemRICH) plot of the significant features from ANOVA **G)** ChemRICH plot of the significant features from C) with the chemical classes in blue are down-regulated and those in red are up-regulated. **H)** ChemRICH plot of the significant features from D). **I)** ChemRICH plot of the significant features from E). **J)** The overlapping extracted ion chromatograms from 6,8-Di-C-ß-D-arabinopyranosylapigenin which showed high fold change and significance in E). **K)** Mirror plot of the experimental $MS^2$ and the $MS^2$ from the spectral library match of the feature from J).

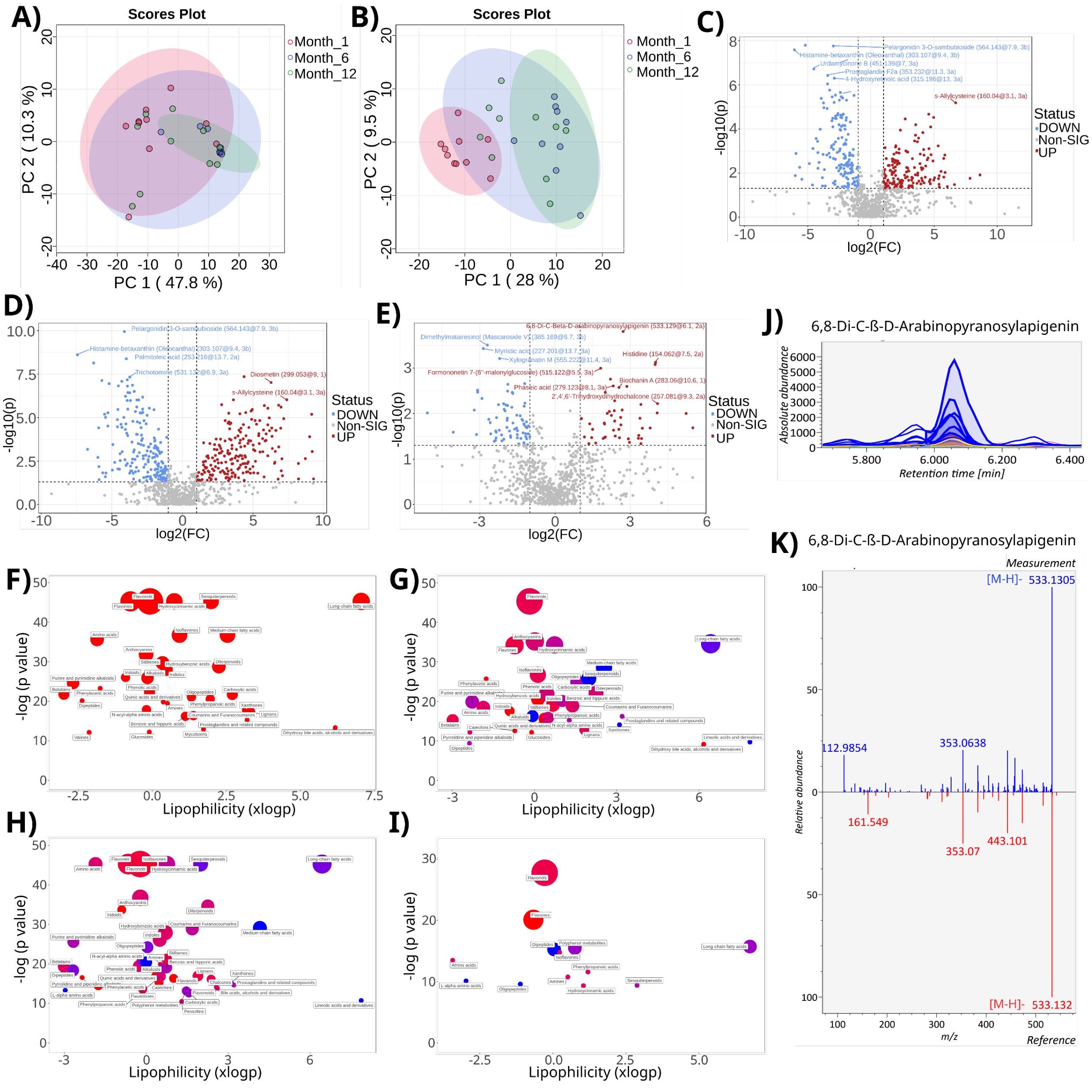

**12. Figure B.7.** Correlations between features in the stool sample and the gut microbiome.

**A)** Spearman rank correlation (p.adj < 0.05) heatmap between the microbes detected by 16S rRNA gene amplicon sequencing (with their species and ASV identifier given) and the features that showed significance by ANOVA (with their m/z, retention time, and identification level given). **B)** A network of the Spearman rank correlation results with microbes (with their genus-level classification and ASV identifier given) colored in green and features (with their m/z, retention time, and identification level given) in orange.

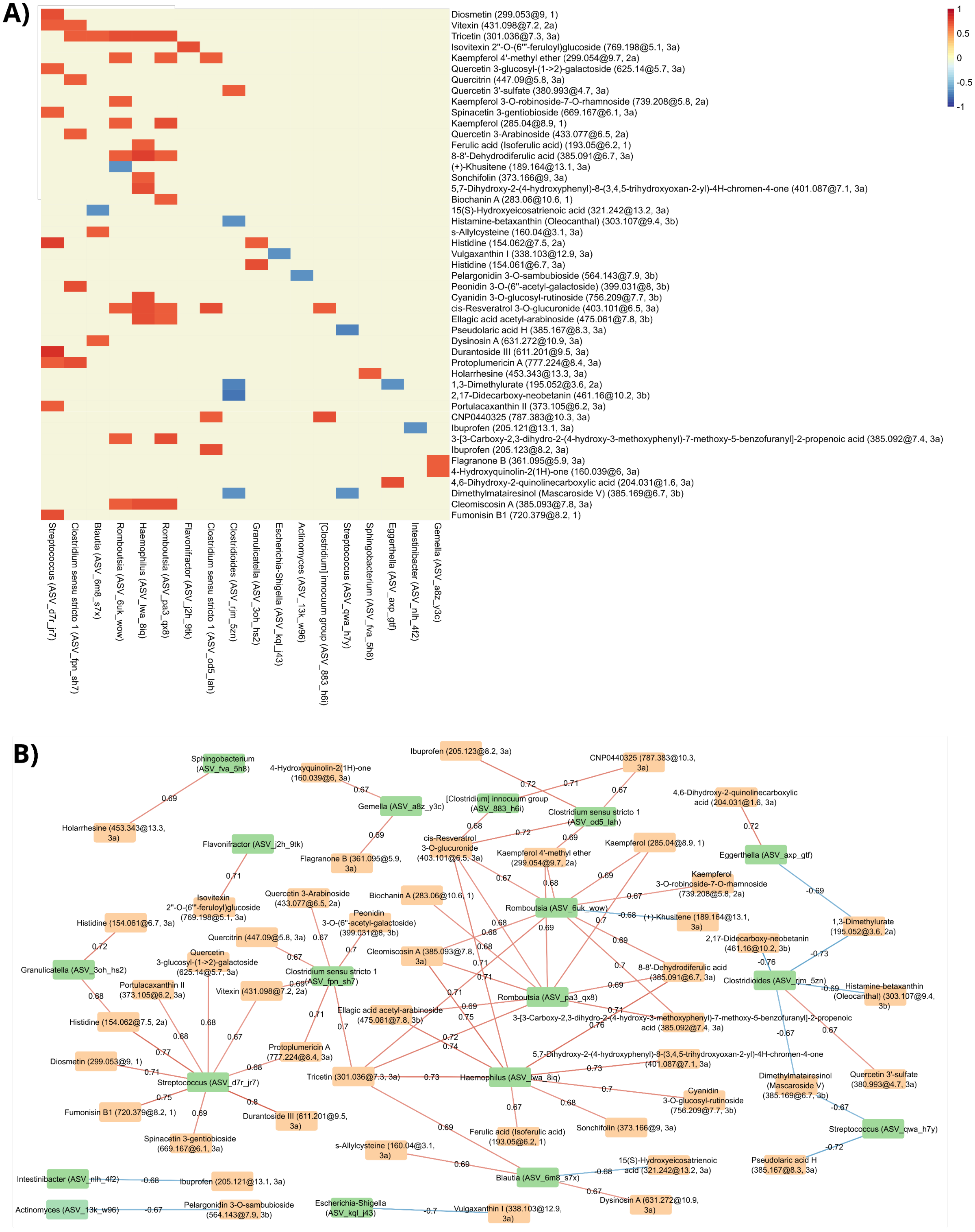
